# CCDC66 couples actin-microtubule crosslinking to KANK1-associated microtubule targeting and focal adhesion turnover

**DOI:** 10.64898/2026.05.19.723697

**Authors:** F. Basak Turan, Ilgin Sezer, Jovana Deretic, Mehmet Gönen, Elif Nur Firat-Karalar

**Author notes:** Corresponding Author: Elif Nur Firat-Karalar, Koç University, Sarıyer, Istanbul, Turkey.

## Abstract

Cell migration requires actin-based protrusion to be coordinated with microtubule (MT)-dependent focal adhesion (FA) remodeling. Although KANK proteins link talin-containing FAs to cortical MT-capture machinery, how this machinery couples to force-bearing actin networks remains unclear. Here, we identify the ciliopathy-associated protein CCDC66 as an actin-MT crosslinker required for FA turnover and migration. CCDC66 depletion impaired collective, Transwell, and random migration, whereas CCDC66 overexpression enhanced wound closure and Matrigel invasion. Mechanistically, CCDC66 loss reduced MT targeting to peripheral FAs and shifted cells into a ROCK-dependent contractile state marked by impaired lamellipodial protrusion, stress fiber accumulation, enlarged long-lived adhesions, and increased RhoA-ROCK signaling. ROCK or formin inhibition suppressed this state, whereas Rac1 activation failed to restore productive protrusion, indicating that CCDC66 maintains the balance between protrusive and contractile actin organization. *In vitro* TIRF reconstitution demonstrated that purified CCDC66 directly crosslinks actin filaments and MTs. In cells, CCDC66 associated with KANK1, supported its peri-adhesion organization, and functioned with KANK1 in an overlapping migration pathway. TCGA analyses further revealed that a coordinated CCDC66/KANK/ROCK-associated module, but not CCDC66 expression alone, stratified outcome in chromophobe renal cell carcinoma. These findings reveal a non-ciliary role for CCDC66 in coupling actin–MT integration to FA turnover and contractile-state control, with relevance to developmental disease and cancer-associated motility.

## Introduction

Cell migration is a fundamental biological process required for development, tissue homeostasis, and wound healing, and its dysregulation contributes to human pathologies, including neurodevelopmental disorders and cancer metastasis. Productive migration depends on polarized cycles of actin-driven protrusion at the leading edge, coupled to contraction of the cell body and retraction at the trailing edge [1, 2]. A key component of this cycle is the dynamic formation and turnover of focal adhesions (FAs), which mechanically couple the actin cytoskeleton to the extracellular matrix and transmit traction forces during migration [3]. The spatiotemporal asymmetry of the actin-based structures in migrating cells is governed by Rho family GTPases [4-6]. At the leading edge, Rac1 promotes branched actin assembly and lamellipodia formation, a process linked to Focal Adhesion Kinase activation and rapid turnover of small, transient adhesions [7]. In contrast, RhoA signaling predominates at the trailing edge and within the cell body. Through ROCK1/2, RhoA promotes myosin light chain (MLC) phosphorylation, thereby driving actomyosin contractility, focal adhesion maturation, and rear retraction.[8, 9]. When this balance is perturbed, productive migration is compromised. Excessive RhoA-ROCK-dependent contractility can promote stress fiber assembly and the maturation or persistence of focal adhesions, thereby slowing adhesion turnover, whereas impaired protrusion dynamics can prevent effective leading-edge advance.

While actin dynamics generate the forces that drive migration, the microtubule (MT) cytoskeleton provides a critical regulatory system that coordinates protrusion, adhesion turnover, and contractility [10, 11]. Polarized MT growth toward the leading edge supports vesicular trafficking and spatial delivery of signaling factors that shape protrusion and adhesion behavior [12, 13]. One way that MTs exert these effects is through their interaction with actin and FAs. Actin stress fibers guide MT growth toward the cell periphery, where MT plus ends target adhesion-proximal regions and promote FA turnover. In this context, MTs locally modulate RhoA-dependent contractility, at least in part through regulation of the MT-associated RhoGEF GEF-H1 [14, 15]. Consequently, defects in MT targeting stabilize adhesions, increase actomyosin-based contractility, and impair migration.

Actin-MT coordination at adhesion sites is mediated by proteins that guide MT plus ends along actin-rich structures, couple MTs to FA-associated capture machinery, and regulate FA turnover and signaling [16-18]. For example, the spectraplakin MACF1/ACF7 guides growing, EB3-positive MTs along actin bundles toward FAs and promotes FA turnover and directional migration [19, 20]. In parallel, EB2-dependent targeting of MAP4K4 to FAs links MT plus-end dynamics to FA disassembly [21]. KANK family proteins provide another major mechanism by coupling FAs to cortical MT capture sites [22, 23]. In particular, KANK1 binds talin at the FA rim and recruits cortical MT-stabilizing complexes (CMSCs) containing LL5β, liprins, CLASPs, and KIF21A, which tether and spatially constrain MT plus ends near adhesions during migration [24-27]. KANK2 also localizes to FA rims and other integrin-dependent adhesion structures and has been linked to adhesion-MT coupling, RhoA regulation, and migration in melanoma and podocyte models, highlighting context-dependent roles of KANK paralogs [28-32]. Although these studies explain how MTs are guided and captured at adhesion sites, how KANK-associated MT capture is physically integrated with force-bearing actin networks and coordinated with local RhoA control during FA turnover remains unresolved.

One candidate for mediating this cytoskeletal integration is CCDC66 (coiled-coil domain containing 66), a MT-associated protein previously characterized in primary cilium biogenesis, ciliary signaling and cell division, where it promotes MT stabilization, organization, and polymerization [33-36]. Consistent with these roles, purified CCDC66 binds MTs and promotes their bundling and stabilization [33, 34, 37]. More recently, we showed that CCDC66 also associates with F-actin and contributes to actin-dependent vesicular trafficking and ectocytosis during cilium disassembly [35]. These findings raise the possibility that CCDC66 coordinates actin and MT networks, but whether it directly crosslinks actin filaments and MTs, and whether this activity functions in dynamic processes such as FA remodeling and cell migration, remains unknown. Addressing this question is disease-relevant because CCDC66 is linked to retinal degeneration in canine and murine models and has been identified within a Joubert syndrome-associated ciliary tip module, suggesting that CCDC66-associated pathology may involve cytoskeletal defects in addition to ciliary dysfunction [36, 38-40].

Here, we investigated whether CCDC66 coordinates actin-MT crosstalk during during FA remodeling and cell migration. We identify CCDC66 as an actin-MT crosslinking factor that links KANK1-associated MT targeting to FA turnover. CCDC66 depletion disrupts this coordination, reducing MT targeting to FAs and shifting cells into a ROCK-dependent contractile state marked by impaired lamellipodial protrusion, stress fiber accumulation, and enlarged, long-lived FAs. In contrast, CCDC66 overexpression enhances wound closure and Matrigel invasion. Together, our findings reveal a non-ciliary role for CCDC66 in coupling actin-MT integration to FA remodeling and migration, and identify a CCDC66/KANK/ROCK-associated cytoskeletal module with outcome relevance in chromophobe renal cell carcinoma.

## Results

### CCDC66 is required for collective, directed, and random cell migration

To define the cellular functions of CCDC66 in migration, we performed siRNA-mediated loss-of-function experiments in human U2OS osteosarcoma cells, an established model for quantitative analysis of cytoskeletal organization, focal adhesion dynamics, and cell migration [41, 42]. qRT-PCR confirmed ∼80% CCDC66 mRNA knockdown using a previously validated siRNA (Fig. S1A) [33]. We first examined collective migration using wound-healing assays. Relative to controls, CCDC66 depletion reduced wound closure by 76%, with depleted cells reaching only 12% closure by 24 h (Fig. 1A,B). Defective wound closure was also observed in hTERT-RPE1 cells, indicating that the requirement for CCDC66 in collective migration is not restricted to U2OS cells (Fig. S2A).

**Figure 1.**
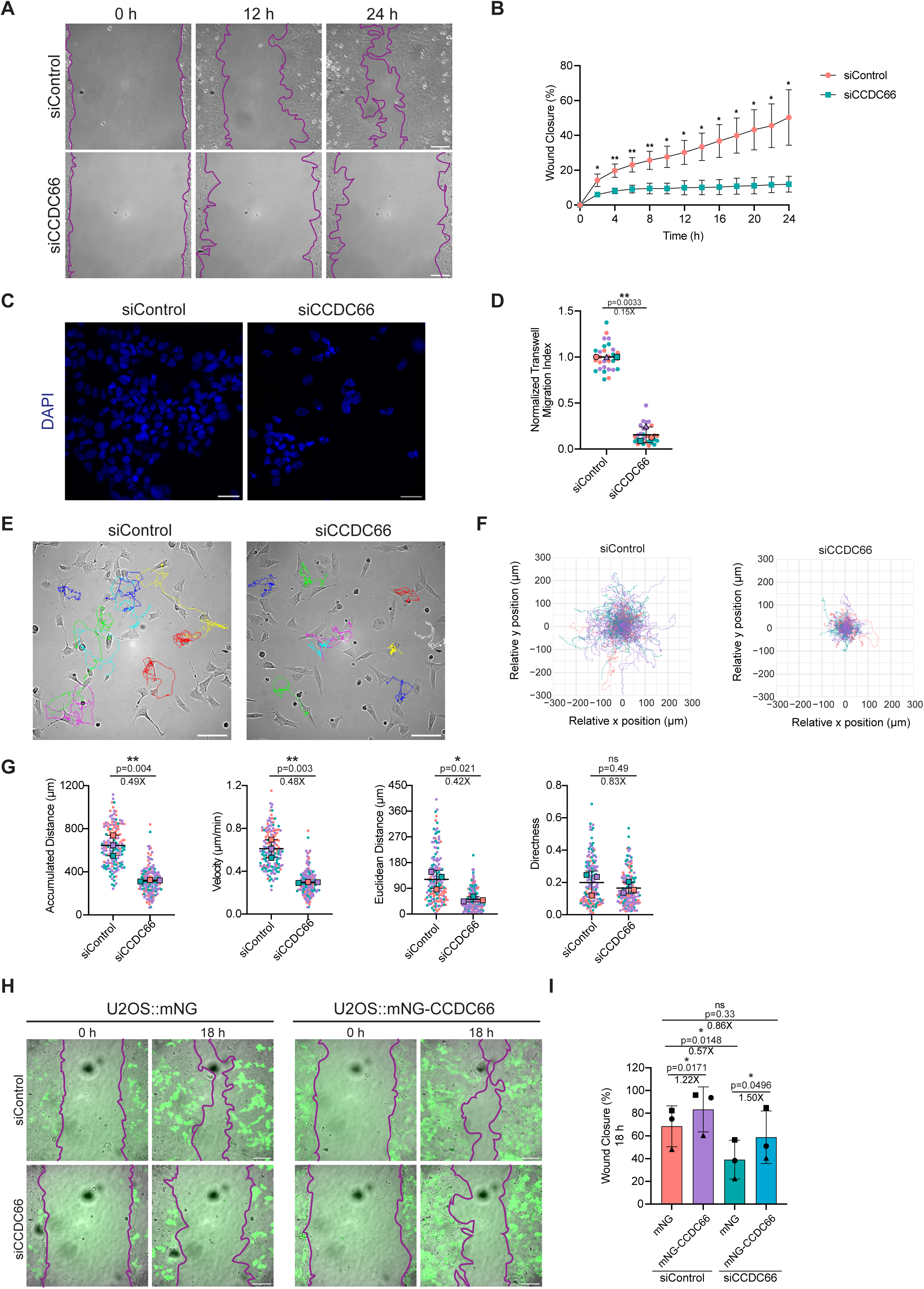
CCDC66 is required for collective, directed, and random cell migration. **(A)** Representative phase-contrast images of a wound healing assay in U2OS cells transfected with control (siControl) or CCDC66-targeting (siCCDC66) siRNAs at 0, 12, and 24 h post-wound. Purple lines outline the migrating wound edges. Scale bars, 100 µm. **(B)** Quantification of wound closure (%) shown in (A) over a 24-h period. Wound closure was calculated as the percentage reduction in wound area relative to the initial wound area at 0 h. Data are presented as mean ± S.E.M. from *n = 3* independent experiments. Asterisks indicate statistical significance at the corresponding time points (*p < 0.05, **p < 0.01; paired Student’s *t* test). **(C)** Transwell migration assay with U2OS transfected with control or CCDC66 siRNAs Representative images of DAPI-stained nuclei of U2OS cells that migrated to the lower surface of a Transwell membrane after 48 h. Scale bars, 40 µm. **(D)** Quantification of Transwell migration, presented as the number of DAPI-positive cells on the lower surface of the membrane and normalized to the control condition. Colored dots represent individual measurements, with each color corresponding to one independent experiment. Data represent mean ± S.E.M. from *n = 3* independent experiments. Fold changes relative to siControl are indicated above the graphs. Asterisks denote statistical significance (**p < 0.01; paired Student’s *t* test). **(E)** Time-lapse imaging assay performed to monitor the random cell migration dynamics. Representative phase-contrast images of U2OS cells, transfected with control or CCDC66 siRNAs, are shown overlaid with individual cell tracks representing their cumulative migratory trajectories over the entire 16 h period. Images were captured every 6 min. Scale bars, 100 µm. **(F)** Trajectory (rose) plots showing the migration paths of all tracked cells, normalized to the origin (0,0). Each color corresponds to one independent experiment. **(G)** Quantification of single-cell random migration parameters of accumulated distance, velocity, Euclidean distance, and directness. Directness was calculated as the ratio of Euclidean distance to accumulated distance. Colored dots represent individual tracked cells, with each color corresponding to one independent experiment; statistical analysis was performed on biological replicate means. Data represent mean ± S.E.M. from *n = 3* independent experiments. Fold changes relative to siControl are indicated above the graphs. Asterisks denote statistical significance (**p < 0.01, *p < 0.05, ns: not significant; paired Student’s *t* test). **(H)** Representative phase-contrast and fluorescence overlay images of a wound healing rescue assay in U2OS cells. Doxycycline-inducible U2OS cell lines expressing either mNG or siRNA-resistant mNG-CCDC66 were transfected with control or CCDC66 siRNAs. Confluent cell monolayers were scratched, and images were acquired at 0 and 18 hours post-scratch are shown. Purple lines outline the migrating wound edges, and green fluorescence confirms the expression of the mNG-tagged constructs. Scale bars, 100 µm. **(I)** Quantification of the wound healing percentage from the assay shown in (H). Wound closure was calculated as the percentage reduction in wound area relative to the initial wound area at 0 h. Data are presented as mean ± S.E.M. from *n = 3* independent experiments. Symbols represent independent experiments. Fold changes and exact p-values are indicated above the graph. Asterisks denote statistical significance (*p < 0.05, ns: not significant; one-way ANOVA with Tukey’s post hoc test).

To determine whether the wound-healing defect reflected impaired migration rather than a secondary consequence of CCDC66’s known roles in cell division [33], we repeated the assay in the presence of the DNA synthesis inhibitor mitomycin C. Under these growth-inhibited conditions, the phenotype persisted, with CCDC66 depletion reducing wound closure by 47% relative to mitomycin-treated controls (Fig. S1B, C). In addition, centrosome/MTOC reorientation toward the wound edge was not altered (Fig. S1D,E), suggesting that global front-rear polarity remains intact despite CCDC66 loss. Having excluded major contributions from proliferation and polarity defects, we next asked whether CCDC66 is required for directional single-cell migration. In Transwell assays, far fewer CCDC66-depleted cells traversed the membrane than control cells (Fig. 1C), corresponding to an 84% reduction in migrated cells per field (Fig. 1D).

We then asked whether CCDC66 is also required for cell movement in the absence of external directional cues. To this end, we tracked individual U2OS cells over 16 h and quantified random migration behavior (Fig. 1E-G). CCDC66 depletion restricted single-cell movement, as evident from the more compact trajectory plots and the reduced spread of rose plots around the origin (Fig. 1E,F). Quantitative analysis confirmed that accumulated distance, velocity, and Euclidean distance were each reduced by approximately 50%, whereas directness, measured as the persistence index, remained unchanged at ∼0.2 in both conditions (Fig. 1G). Together, these data indicate that CCDC66 is required for the speed and efficiency of single-cell movement, rather than for intrinsic directional persistence under random migration conditions.

To assess whether these migration defects were specific to loss of CCDC66, we performed wound-healing assays using doxycycline-inducible U2OS cells expressing mNeonGreen (mNG) or siRNA-resistant mNG-CCDC66 (Fig. 1H,I). Validation of these lines by co-staining with the centriolar satellite marker PCM1 and centrosome marker gamma-tubulin confirmed the expected localization of mNG and the centrosomal and satellite localization of mNG-CCDC66 (Fig. S1F). Re-expression of CCDC66 in CCDC66-depleted cells increased wound closure relative to the mNG control, consistent with partial restoration of migration. In parallel, induction of mNG-CCDC66 in control cells further increased wound closure, indicating that CCDC66 overexpression promotes migration. Consistent with this gain-of-function effect, mNG-CCDC66-expressing cells also showed increased invasion through Matrigel-coated Transwell filters relative to mNG-expressing control cells (Fig. S1G,H). Together, these loss-, rescue-, and gain-of-function experiments support CCDC66 as a positive regulator of cell motility.

### CCDC66 depletion impairs lamellipodial protrusion and leads to stress fiber and focal adhesion accumulation

Efficient cell migration requires a spatial balance between Rac-dependent leading-edge protrusions (lamellipodia) and Rho-dependent contractile actomyosin structures (stress fibers and focal adhesions) within the cell body and trailing edge [3, 9]. To determine whether CCDC66 regulates this organization, we first visualized the actin cytoskeleton in cells expressing RFP-LifeAct. While control cells displayed broad lamellipodia and thin stress fibers, CCDC66-depleted cells had prominent bundled stress fibers and reduced broad protrusive edges both in subconfluent cultures and at wound edges, consistent with increased contractile organization (Fig. 2A, Fig. S2A). To quantify this shift in actin organization, we examined endogenous F-actin networks by phalloidin staining together with the boundary marker β-catenin (Fig. 2B). This analysis confirmed significant increases in both stress fiber number (∼2.1-fold) and stress fiber thickness (∼1.3-fold) following CCDC66 depletion (Fig. 2C).

**Figure 2.**
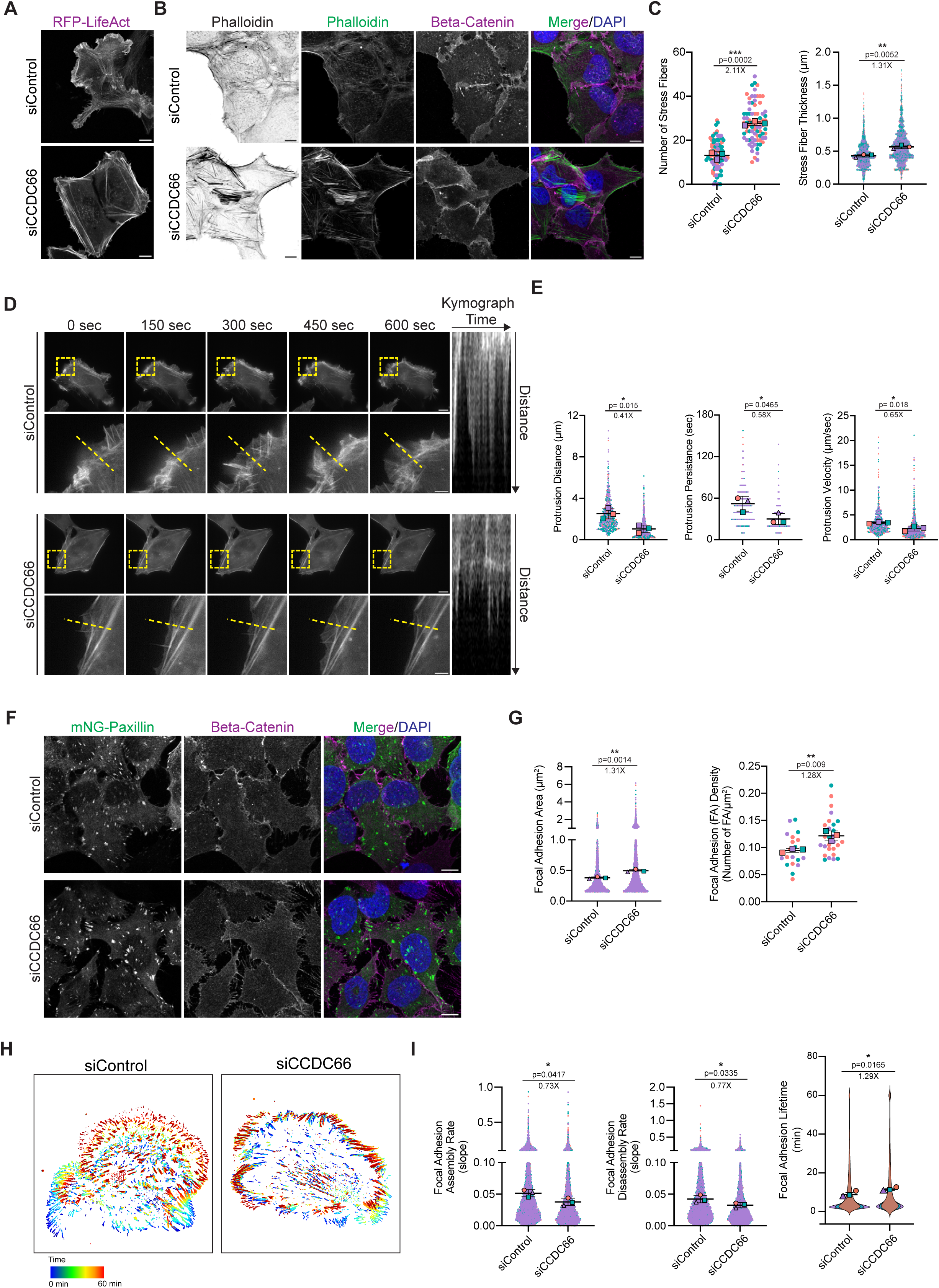
CCDC66 is required for proper actin organization and the regulation of focal adhesion size, density, and dynamics. **(A)** Representative images of U2OS cells expressing RFP-LifeAct (shown in grayscale), transfected with control or CCDC66 siRNAs, to visualize actin cytoskeleton. Scale bars, 10 µm. **(B)** Representative immunofluorescence images of U2OS cells transfected with control or CCDC66 siRNAs. Cells were stained with phalloidin to visualize F-actin (green) and co-stained for the cell boundary marker β-catenin (magenta). Nuclei were counterstained with DAPI (blue). Scale bars, 10 µm. **(C)** Quantification of stress fiber number per cell (left) and stress fiber thickness (µm) (right) based on the phalloidin-stained cells shown in (B). Data represent mean ± S.E.M. from *n = 3* independent experiments. Colored dots represent individual measurements, with each color corresponding to one independent experiment. Fold changes relative to siControl are indicated above the graphs. Asterisks denote statistical significance (**p < 0.01, ***p < 0.001; paired Student’s *t* test). **(D)** Live imaging of U2OS cells stably expressing RFP-LifeAct (shown in grayscale) and transfected with control or CCDC66 siRNAs. Cells were imaged every 10 s for 10 min. Yellow dashed boxes indicate the regions shown at higher magnification below, and yellow dashed lines indicate the axis used to generate the corresponding kymographs (right) for protrusion analysis. Scale bars, 10 µm (upper panels) and 3 µm (lower panels). **(E)** Quantification of protrusion dynamics from the kymographs in (D), including protrusion distance (µm), protrusion persistence (sec), and protrusion velocity (µm/sec). Data represent mean ± S.E.M. from *n = 3* independent experiments. Colored dots represent individual measurements, with each color corresponding to one independent experiment. Fold changes relative to siControl are indicated above the graphs. Asterisks denote statistical significance (*p < 0.05; paired Student’s *t* test). **(F)** Representative images of fixed U2OS cells expressing mNG-Paxillin (green), transfected with control or CCDC66 siRNAs, to visualize focal adhesions. Cells were co-stained for β-catenin (magenta) and DAPI (blue). Scale bars, 10 µm. **(G)** Quantification of focal adhesion morphology based on the mNG-Paxillin signal shown in (F), including focal adhesion area (µm²), and focal adhesion density (number of FAs per µm² of cell area). Data represent mean ± S.E.M. from *n = 3* independent experiments. Colored dots represent individual measurements, with each color corresponding to one independent experiment. Fold changes relative to siControl are indicated above the graphs. Asterisks denote statistical significance (**p < 0.01; paired Student’s *t* test). **(H)** Representative temporal color-coded projections derived from time-lapse imaging of U2OS cells expressing mNG-Paxillin and transfected with control or CCDC66 siRNAs. Images were acquired every 1 min for 60 min. The color bar indicates time, ranging from 0 min (blue) to 60 min (red). **(I)** Quantification of focal adhesion dynamics, specifically Focal Adhesion Assembly and Disassembly Rates, derived from the time-lapse sequences shown in (H). Rates were calculated from changes in focal adhesion area over time in mNG-Paxillin time-lapse sequences. Data represent mean ± S.E.M. from *n = 3* independent experiments. Colored dots represent individual measurements, with each color corresponding to one independent experiment. Fold changes relative to siControl are indicated above the graphs. Asterisks denote statistical significance (*p < 0.05; paired Student’s *t* test).

We next asked whether this altered actin architecture affected leading-edge dynamics. Live imaging of RFP-LifeAct-expressing cells followed by kymograph analysis showed smooth, persistent edge advance in control cells, whereas CCDC66-depleted cells exhibited frequent, short-lived protrusion-retraction cycles with little net edge displacement (Fig. 2D). Quantification revealed significant reductions in protrusion distance (∼59%), protrusion persistence (∼42%), and protrusion velocity (∼35%), indicating that CCDC66 is required to sustain productive lamellipodial protrusion (Fig. 2E).

Because stress fibers terminate at FAs and transmit contractile forces to the extracellular matrix [43], we next examined whether the altered actin architecture in CCDC66-depleted cells was associated with changes in adhesion morphology. CCDC66-depleted U2OS cells accumulated enlarged, elongated mNG-Paxillin-positive adhesions, and quantification confirmed significant increases in both FA area and FA density (Fig. 2F,G). A similar actin and adhesion phenotype was observed in RPE1 cells, where CCDC66 depletion induced prominent phalloidin-positive stress fibers and enlarged paxillin-positive adhesions (Fig. S2B,C). To test whether the FA phenotype was specific to CCDC66 loss, we analyzed U2OS rescue lines constitutively expressing either mNG or siRNA-resistant mNG-CCDC66. CCDC66 expression reduced the increased FA area and density observed after CCDC66 depletion toward control levels, whereas mNG alone did not (Fig. S2D,E).

Because steady-state FA size and abundance reflect the balance between adhesion formation, maturation, and disassembly [44], we next asked whether CCDC66 depletion alters FA turnover. To test this, we performed time-lapse imaging of mNG-Paxillin (Fig. 2H). Temporal color-coding of adhesion tracks over a 60-min period showed that FAs turned over dynamically near the leading edge in control cells, whereas adhesions in CCDC66-depleted cells persisted for longer periods (Fig. 2H). Quantification revealed significant decreases in both FA assembly and disassembly rates, together with a significant increase in FA lifetime in CCDC66-depleted cells (Fig. 2I). Together, these kinetic data indicate that loss of CCDC66 broadly slows FA turnover, resulting in longer-lived adhesions that parallel the observed increases in FA size and density.

Collectively, these results show that CCDC66 is required to maintain the balance between protrusive and contractile actin organization. Its depletion shifts cells toward a stress fiber-rich state characterized by reduced lamellipodial productivity, bundled stress fibers, and enlarged, long-lived FAs with reduced turnover.

### CCDC66 depletion shifts the balance between branched and linear actin assembly pathways

The concomitant reduction in lamellipodial protrusion and accumulation of bundled stress fibers suggested that CCDC66 depletion alters the balance between branched and linear actin assembly pathways. To test this, we pharmacologically perturbed formin- and Arp2/3-dependent actin assembly and assessed their contributions to the CCDC66-depletion phenotype (Fig. S3).

Treatment with the formin inhibitor SMIFH2 reduced FA area, FA density, and stress fiber number in control and CCDC66-depleted cells, consistent with the established role of formins in the assembly of stress fibers and the maturation of focal adhesions (Fig. S3A-D) [45]. In CCDC66-depleted cells, these parameters were restored to the levels observed in DMSO-treated controls, indicating a significant reversal of the excessive stress fiber and focal adhesion accumulation. Notably, this structural rescue was accompanied by improved migration, yielding a ∼50% relative increase in wound closure (Fig. S3E,F). These results indicate that the stress fiber-rich morphology with enlarged FAs contributes to the migration defect caused by CCDC66 depletion.

In contrast, inhibition of Arp2/3 with CK-666 strongly reduced wound closure in control cells by ∼60% (Fig. S3G,H), consistent with the requirement for branched actin assembly during productive migration. In CCDC66-depleted cells, where basal migration was already impaired, CK-666 produced a smaller additional decrease by ∼40%. This suggests that CCDC66 loss already compromises Arp2/3-dependent protrusive migration, while leaving a limited residual contribution to motility. Together with the SMIFH2-mediated reduction in stress fibers and FAs, these data support a model in which CCDC66 depletion biases actin organization toward formin-dependent linear actin structures and away from productive Arp2/3-dependent protrusion.

### CCDC66 depletion induces a ROCK-dependent contractile state that impairs cell migration

Given the stress fiber accumulation and FA enlargement observed upon CCDC66 depletion, we next asked whether RhoA-ROCK signaling was altered. Biochemical pulldown assays using RBD beads showed increased levels of GTP-bound mNG-RhoA following CCDC66 depletion (Fig. 3A). Consistent with elevated RhoA-ROCK pathway activity, immunoblotting and immunofluorescence analysis showed increased phosphorylation of myosin regulatory light chain (p-MLC), a downstream marker of actomyosin contractility (Fig. 3B, C).

**Figure 3.**
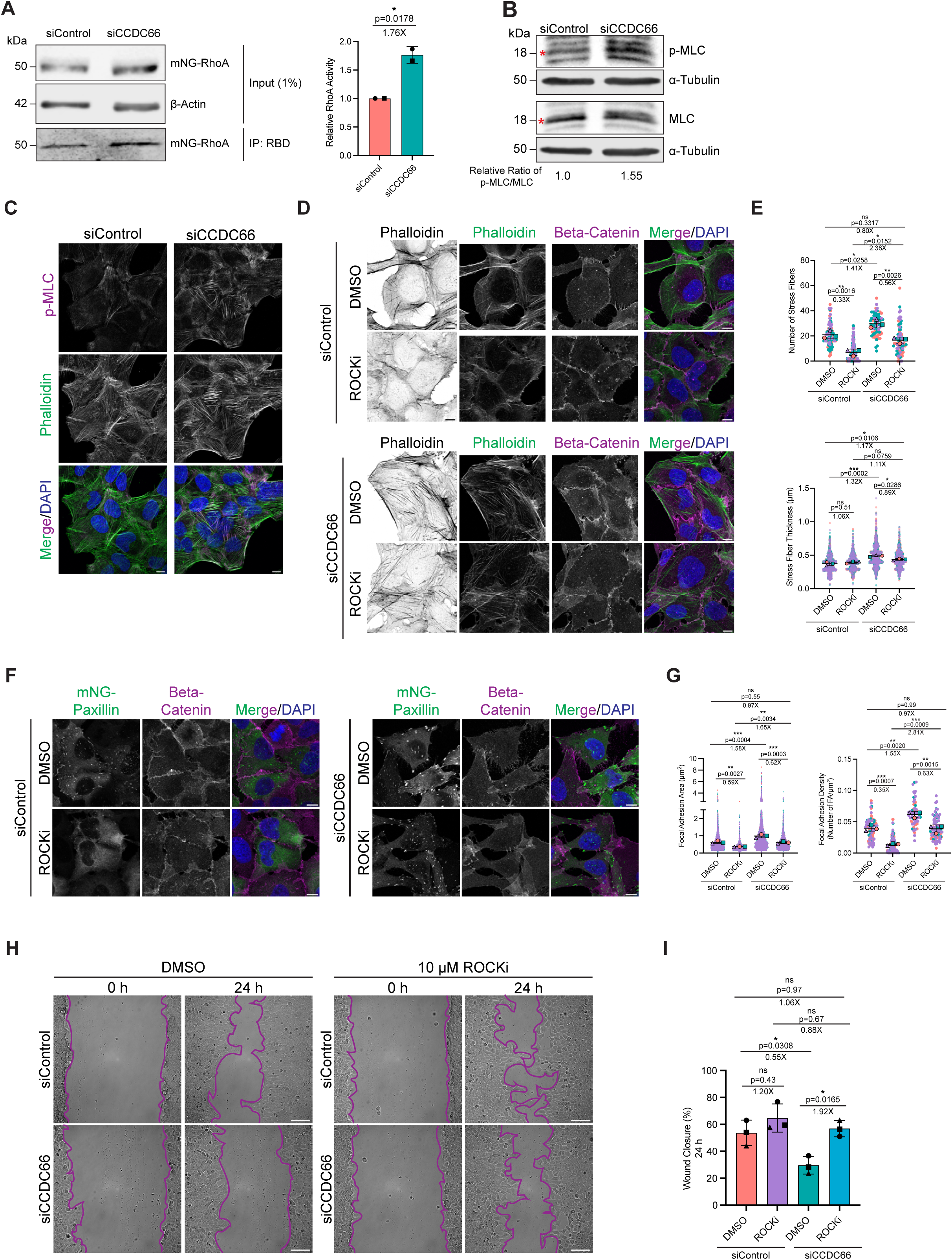
RhoA-ROCK signaling contributes to stress fiber accumulation, focal adhesion enlargement, and migration defects in CCDC66-depleted cells. **(A)** Active RhoA pulldown assay in U2OS cells expressing mNG-RhoA and transfected with control or CCDC66 siRNAs. Equal amounts of total cell lysates were incubated with RBD beads to precipitate GTP-bound mNG-RhoA. Representative immunoblots show total mNG-RhoA in the input and active mNG-RhoA in the RBD pulldown. β-actin was used as a loading control. Quantification of relative RhoA activity is shown on the right. Relative activity was calculated as the ratio of RBD-bound mNG-RhoA to input mNG-RhoA normalized to β-actin, and values were normalized to siControl. Data represent mean ± S.E.M. from *n = 2* independent experiments. Symbols represent independent experiments. Fold change relative to siControl is indicated above the graph. Asterisks denote statistical significance (*p < 0.05; paired Student’s *t* test). **(B)** Immunoblot analysis of phosphorylated myosin light chain (p-MLC) and total myosin light chain (MLC) in whole-cell lysates from U2OS cells transfected with control or CCDC66 siRNAs. α-tubulin was used as a loading control. Red markers indicate the bands used for quantification. Quantification of the p-MLC/total MLC ratio is shown below the blots. p-MLC and total MLC signals were normalized to their respective α-tubulin loading controls, and the normalized p-MLC value was divided by the normalized total MLC value. Values were normalized to siControl. Quantification represents *n = 1* experiment and is shown as a representative measurement. **(C)** Representative immunofluorescence images of U2OS cells transfected with control or CCDC66 siRNAs and stained for p-MLC (magenta), F-actin using phalloidin (green), and DAPI (blue). Scale bars, 10 µm. **(D)** Representative immunofluorescence images of U2OS cells transfected with control or CCDC66 siRNAs and treated with DMSO or the ROCK inhibitor Y-27632 (ROCKi, 10 µM) for 2 h before fixation. Cells were stained with phalloidin to visualize F-actin (green), co-stained for the cell boundary marker β-catenin (magenta), and counterstained with DAPI (blue). Scale bars, 10 µm. **(E)** Quantification of stress fiber number per cell and stress fiber thickness (µm) based on the phalloidin-stained cells shown in (D). Data represent mean ± S.E.M. from *n = 3* independent experiments. Colored dots represent individual measurements, with each color corresponding to one independent experiment. Fold changes and exact p values are indicated above the graphs. Asterisks denote statistical significance (*p < 0.05, **p < 0.01, ***p < 0.001, ns = not significant; two-way ANOVA with Tukey’s post hoc test). **(F)** Representative immunofluorescence images of U2OS cells stably expressing mNG-Paxillin (green), transfected with control or CCDC66 siRNAs, and treated with DMSO or Y-27632 (ROCKi, 10 µM) for 2 h before fixation. Cells were co-stained for β-catenin (magenta) and DAPI (blue). Scale bars, 10 µm. **(G)** Quantification of focal adhesion morphology based on the mNG-Paxillin signal shown in (F), including focal adhesion area (µm²) and focal adhesion density (number of FAs per µm² of cell area). Data represent mean ± S.E.M. from *n = 3* independent experiments. Colored dots represent individual measurements, with each color corresponding to one independent experiment. Fold changes and exact p values are indicated above the graphs. Asterisks denote statistical significance (**p < 0.01, ***p < 0.001, ns = not significant; two-way ANOVA with Tukey’s post hoc test). **(H)** Representative phase-contrast images from wound-healing assays in U2OS cells transfected with control or CCDC66 siRNAs and treated with DMSO or Y-27632 (ROCKi, 10 µM). At 48 h post-transfection, confluent monolayers were scratched and imaged at 0 and 24 h post-wounding. Purple lines outline the migrating wound edges. Scale bars, 100 µm. **(I)** Quantification of wound closure from the assay shown in (H). Wound closure was calculated as the percentage reduction in wound area relative to the initial wound area at 0 h. Data are presented as mean ± S.E.M. from *n = 3* independent experiments. Symbols represent independent experiments. Fold changes and exact p values are indicated above the graph. Asterisks denote statistical significance (*p < 0.05, ns = not significant; two-way ANOVA with Tukey’s post hoc test).

To determine whether ROCK activity contributes functionally to the actin and adhesion phenotypes caused by CCDC66 depletion, we treated cells with the ROCK inhibitor Y-27632. ROCK inhibition suppressed the CCDC66-depletion-induced increases in both stress fiber number and stress fiber thickness, restoring them to levels comparable to DMSO-treated control cells (Fig. 3D,E). Similarly, Y-27632 reduced the increased FA area and density observed in CCDC66-depleted cells to the control levels (Fig. 3F,G). We then asked whether the suppression of these stress fiber and FA defects translated to a functional rescue of migration. In wound-healing assays, ROCK inhibition had only a modest, non-significant effect on the wound closure of control cells (1.20-fold increase) (Fig. 3H,I). In contrast, Y-27632 treatment significantly increased wound closure in CCDC66-depleted cells (1.92-fold), restoring migration to levels comparable to control cells (Fig. 3H, I). Together, the increased RhoA-GTP levels and the rescue of stress fiber, FA, and migration defects by Y-27632 indicate that CCDC66 depletion induces a ROCK-dependent hypercontractile state that limits efficient migration.

Because CCDC66-depleted cells showed defective lamellipodial protrusion, we next examined whether enhancing Rac-dependent protrusive signaling could overcome the migration defect. To this end, we generated U2OS cells stably expressing constitutively active Rac1 (mNG-Rac1 Q61L) and analyzed wound closure (Fig. S4A,B). Rac1 Q61L expression produced only a modest increase in wound closure in both control (1.12-fold) and CCDC66-depleted cells (1.20-fold) relative to their corresponding mNG controls. Notably, wound closure in CCDC66-depleted cells expressing Rac1 Q61L was significantly impaired compared with Rac1 Q61L-expressing control cells (Fig. S4B), indicating that Rac1 activation is not sufficient to restore migration upon CCDC66 loss.

To understand why Rac1 activation failed to rescue the migration defect, we examined leading-edge organization and dynamics using time-lapse imaging of RFP-LifeAct (Fig. S4C, D). In control cells, Rac1 Q61L induced dynamic peripheral ruffling and broad lamellipodial extensions, as seen in the time series and temporal color-coded projections (Fig. S4C). In contrast, CCDC66-depleted cells expressing Rac1 Q61L failed to restore broad sheet-like protrusions and retained prominent linear actin bundles extending to the cell periphery. Instead, edge activity was limited to smaller, less productive protrusive events (Fig. S4C). Quantification showed that Rac1 Q61L did not rescue protrusion distance, persistence, or velocity in CCDC66-depleted cells, which remained impaired relative to Rac1 Q61L-expressing control cells (Fig. S4D). Thus, CCDC66 loss is not explained by insufficient Rac activation alone. Instead, CCDC66 depletion induces a ROCK-dependent contractile state that limits productive protrusion and migration.

### CCDC66 directly crosslinks actin and microtubules and promotes microtubule targeting to focal adhesions

To understand how CCDC66 regulates RhoA activity and FA turnover, we focused on actin-microtubule crosstalk, a key mechanism that supports microtubule (MT) targeting to adhesions [18]. During cell migration, dynamic MTs track along stress fibers to reach peripheral FAs, where they promote FA disassembly and turnover, in part through regulation of RhoA signaling [10, 11]. We previously showed that CCDC66 binds and and stabilizes MTs and associates with F-actin, promoting actin bundling [33, 35]. However, whether CCDC66 can directly couple actin and MTs into a single organized network remained unknown. We therefore tested whether purified CCDC66 is sufficient to crosslink F-actin and MTs *in vitro*, and then asked whether CCDC66 supports MT targeting to FAs in cells.

To test the crosslinking activity directly, we performed *in vitro* reconstitution assays using mNG-CCDC66 purified from insect cells as previously described [33]. Using TIRF microscopy, we examined whether mNG-CCDC66 could crosslink soluble rhodamine-labeled F-actin to taxol-stabilized MTs immobilized on the coverslip surface. Control reactions containing mNG alone were performed in parallel. Consistent with its role as a microtubule-associated protein, mNG-CCDC66, but not mNG alone, bound directly to the immobilized microtubules (Fig. 4A). In reactions containing F-actin alone, mNG-CCDC66 organized the actin filaments into prominent bundles, whereas the mNG control exhibited an unorganized actin meshwork. When both polymers were present, mNG-CCDC66 recruited soluble F-actin to immobilized MTs and promoted their co-alignment, forming dense actin-MT bundles (Fig. 4A). In contrast, actin and MTs remained as largely independent networks in the mNG control reactions. Together, these data indicate that CCDC66 is sufficient to crosslink and organize actin and MT networks *in vitro*.

**Figure 4.**
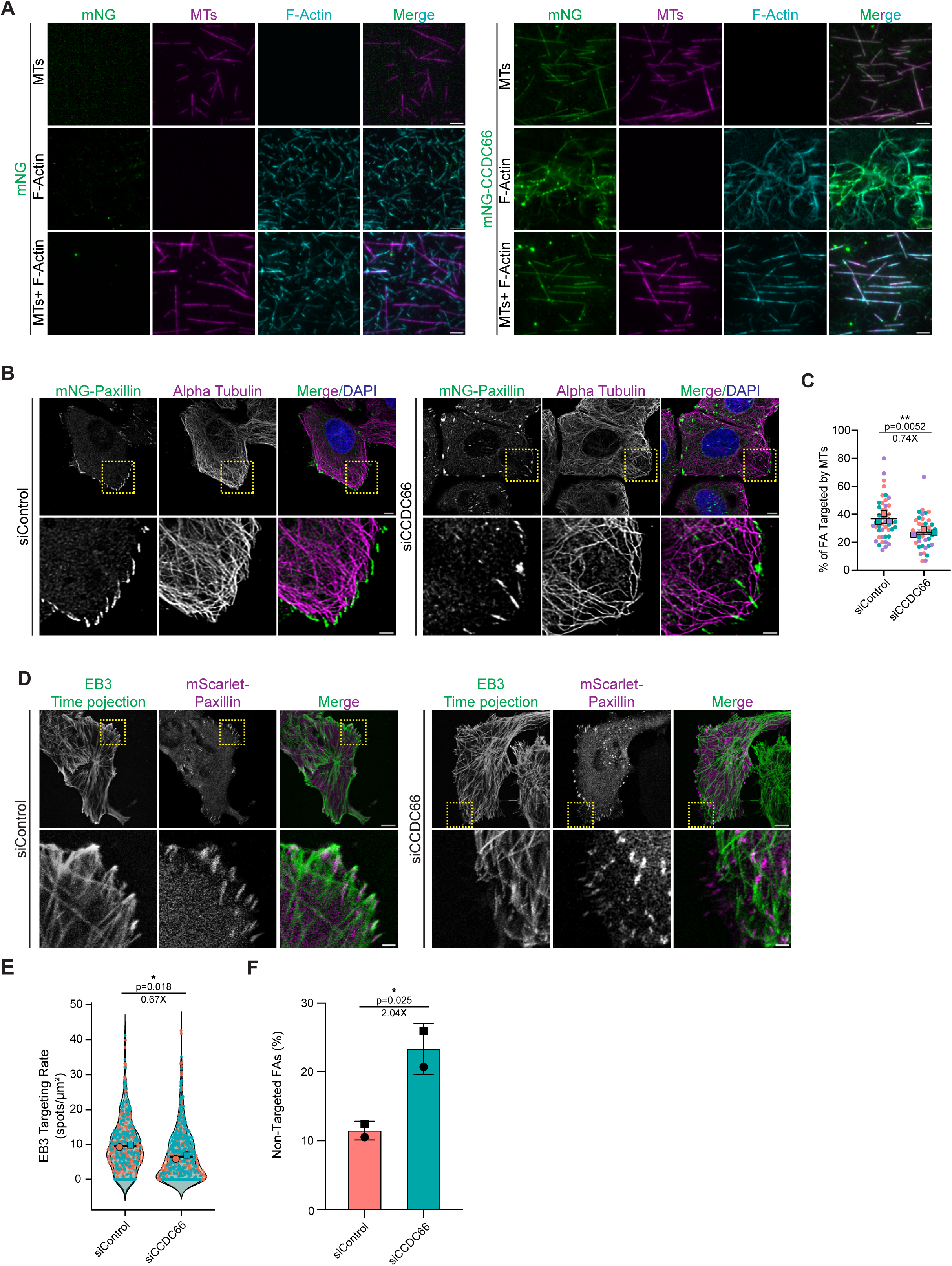
CCDC66 crosslinks actin and microtubules and promotes microtubule targeting to focal adhesions. **(A)** *In vitro* TIRF microscopy analysis of actin-microtubule crosslinking by CCDC66. Assays were performed in flow chambers using silanized coverslips. Where indicated, Alexa Fluor 647-labeled microtubules, stabilized with GMP-CPP and Taxol, were immobilized on the coverslip surface using anti-β-tubulin antibody. His-MBP-mNeonGreen-CCDC66 (mNG-CCDC66, 100 nM) or His-MBP-mNeonGreen control protein (mNG, 100 nM) was incubated with immobilized microtubules (MTs, magenta), soluble rhodamine-labeled F-actin (cyan), or both polymers together. Representative TIRF images show mNG-CCDC66 association with MTs, bundling of F-actin, and co-alignment of F-actin with immobilized MTs when both polymers are present. mNG control reactions are shown in parallel. Scale bars, 3 µm. **(B)** Representative immunofluorescence images of U2OS cells stably expressing mNG-Paxillin (green), transfected with control or CCDC66 siRNAs, and fixed 48 h post-transfection. Cells were stained for α-tubulin to visualize microtubules (magenta), and nuclei were counterstained with DAPI (blue). Yellow dashed boxes indicate the peripheral regions shown at higher magnification below. Scale bars, 5 µm for full-cell images and 2 µm for magnified regions. **(C)** Quantification of microtubule targeting to peripheral focal adhesions from the experiment shown in (B). Peripheral mNG-Paxillin-positive focal adhesions were scored as microtubule-targeted when an α-tubulin-positive microtubule end contacted or terminated at the adhesion. Focal adhesions lacking microtubule contact were scored as non-targeted. Data represent mean ± S.E.M. from *n = 3* independent experiments. Colored dots represent individual measurements, with each color corresponding to one independent experiment. Fold change and exact p value are indicated above the graph. Asterisks denote statistical significance (**p < 0.01; paired Student’s *t* test). **(D)** Live-cell confocal imaging of microtubule plus-end targeting to focal adhesions. U2OS cells expressing GFP-EB3 (green, shown as time projections) and mScarlet-Paxillin (magenta) were transfected with control or CCDC66 siRNAs and imaged 48 h post-transfection. Time-lapse sequences were acquired at 1-s intervals for 1 min. Yellow dashed boxes indicate the peripheral regions shown at higher magnification below. Scale bars, 10 µm for full-cell images and 2 µm for magnified regions. **(E)** Quantification of EB3 targeting rate to focal adhesions from the live-cell imaging shown in (D). EB3 targeting rate was calculated as the number of EB3-positive plus-end targeting events associated with mScarlet-Paxillin-positive focal adhesions, normalized to focal adhesion area. Data are expressed as EB3 spots per µm². Data represent mean ± S.E.M. from *n = 2* independent experiments. Colored dots represent individual measurements, with each color corresponding to one independent experiment. Fold change and exact p value are indicated above the graph. Asterisks denote statistical significance (*p < 0.05; paired Student’s *t* test) **(F)** Percentage of non-targeted focal adhesions from the live-cell imaging shown in (D). Non-targeted focal adhesions were defined as mScarlet-Paxillin-positive adhesions that received no EB3-positive plus-end targeting event during the 1-min imaging period. Data represent mean ± S.E.M. from *n = 2* independent experiments. Dots represent independent experiments. Fold change and exact p value are indicated above the graph. Asterisks denote statistical significance (*p < 0.05; paired Student’s *t* test).

We next tested whether CCDC66 is required for MT targeting to FAs in cells. In fixed U2OS cells expressing mNG-Paxillin, we quantified MT-FA interactions by measuring the fraction of paxillin-positive adhesions contacted by α-tubulin-labeled MT ends (Fig. 4B). In control cells, MT ends extended into the cell periphery and frequently contacted paxillin-positive FAs. CCDC66 depletion reduced the fraction of microtubule-targeted FAs by approximately 26% relative to control cells (Fig. 4C).

To assess dynamic microtubule targeting, we performed live-cell imaging of GFP-EB3 comets together with mScarlet-Paxillin and generated EB3 time projections to visualize cumulative microtubule plus-end trajectories during the imaging period (Fig. 4D). In control cells, EB3-positive plus-end tracks frequently reached peripheral FAs, whereas CCDC66-depleted cells showed fewer EB3 targeting events at adhesions (Fig. 4D, E). Consistently, the fraction of non-targeted FAs was increased approximately twofold following CCDC66 depletion (Fig. 4F). Together, these data indicate that CCDC66 promotes efficient microtubule targeting to FAs, consistent with a role in linking actin-microtubule organization to adhesion turnover.

### CCDC66 interacts with KANK1 and regulates its localization at the focal adhesions

We next asked how CCDC66 is physically linked to adhesion-associated MT capture machinery. To identify candidate factors, we compiled CCDC66 interactors from published proximity-mapping datasets and performed Gene Ontology (GO) enrichment analysis (Fig. 5A) [34, 35, 46]. The enriched categories included cell migration, actin cytoskeleton organization, small GTPase-mediated signaling, and cell adhesion, paralleling the migration, actin, RhoA-ROCK, and FA phenotypes observed upon CCDC66 depletion (Fig. 5A). For example, the network contained small GTPase signaling proteins implicated in FA turnover (e.g. SIPA1L1, ARHGEF6), core FA components (e.g. talin-1 and paxillin), and KANK family proteins. Thus, the proximity network both supported a role for CCDC66 in adhesion-associated cytoskeletal regulation and suggested candidate factors that could link CCDC66 to FAs.

**Figure 5.**
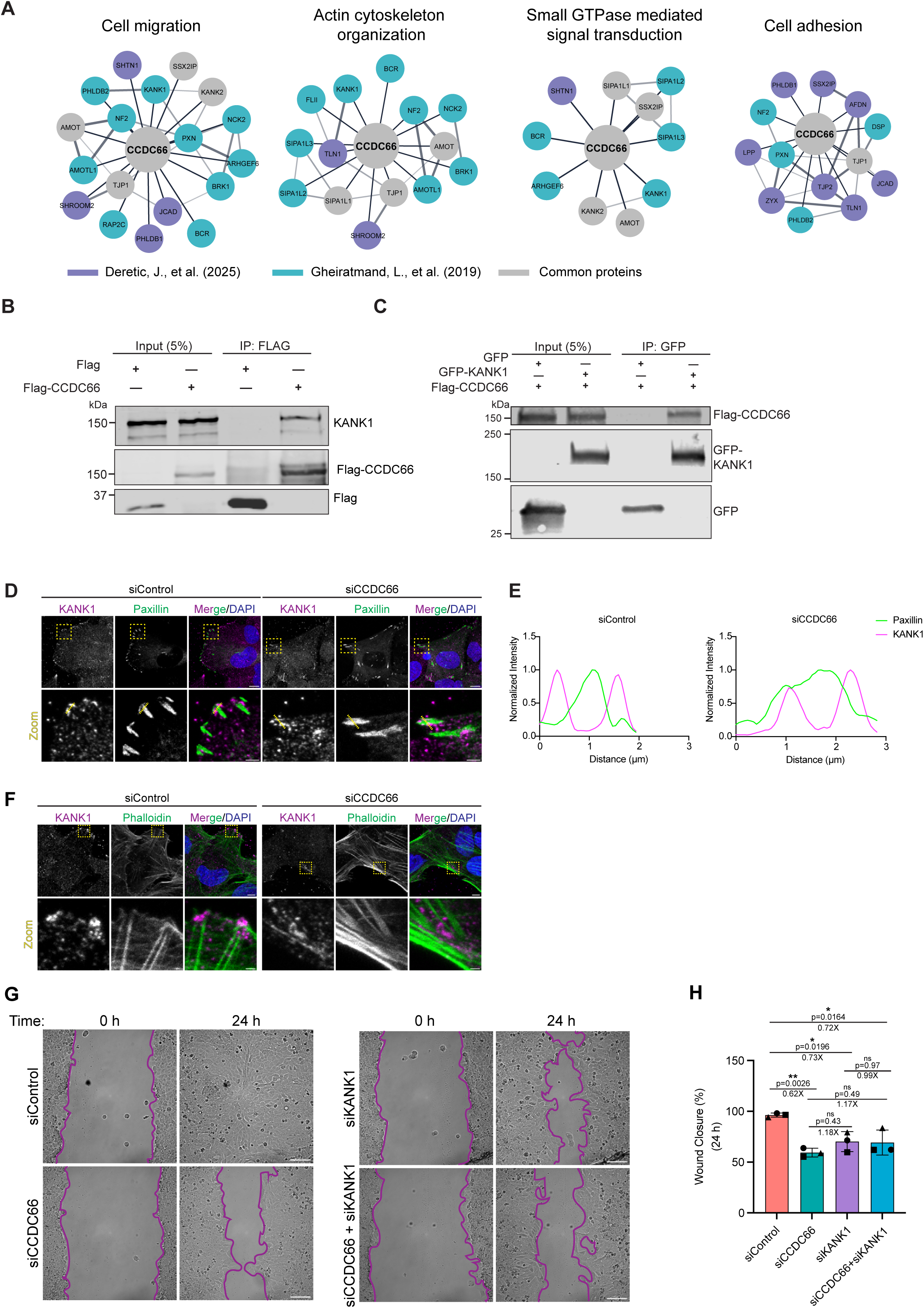
CCDC66 interacts with KANK1, regulates its peri-adhesion organization, and functions with KANK1 during cell migration. **(A)** Bioinformatic analysis of published CCDC66 proximity interactomes. High-confidence interactors (log2(FC) ≥ 1) identified in Gheiratmand et al. (2019) and Deretic et al. (2025) were subjected to Gene Ontology enrichment analysis and visualized as Cytoscape network clusters. Enriched categories included cell migration, actin cytoskeleton organization, small GTPase-mediated signaling, and cell adhesion. Node colors indicate the dataset from which each interactor was identified: Deretic et al. (purple), Gheiratmand et al. (blue), or both datasets (gray). Edge thickness represents STRING interaction confidence scores. **(B)** Co-immunoprecipitation of endogenous KANK1 with FLAG-BirA-CCDC66. HEK293T cells were transfected with FLAG-BirA or FLAG-BirA-CCDC66. At 48 h post-transfection, cell lysates were subjected to anti-FLAG affinity pulldown and analyzed by immunoblotting for endogenous KANK1 and FLAG. FLAG-BirA-CCDC66 co-precipitated endogenous KANK1, whereas FLAG-BirA control did not. Input represents 5% of total cell lysate. **(C)** Co-immunoprecipitation of exogenously expressed CCDC66 and KANK1. HEK293T cells were co-transfected with FLAG-CCDC66 and either GFP alone or GFP-KANK1. At 48 h post-transfection, cell lysates were subjected to GFP-trap pulldown and analyzed by immunoblotting with anti-FLAG and anti-GFP antibodies. FLAG-CCDC66 was co-precipitated with GFP-KANK1 but not with GFP alone. Input represents 5% of total cell lysate. **(D)** Representative immunofluorescence images of U2OS cells stably expressing mNG-Paxillin (green), transfected with control or CCDC66 siRNAs, and stained for endogenous KANK1 (magenta) 48 h post-transfection. Nuclei were counterstained with DAPI (blue). Yellow dashed boxes indicate peripheral regions shown at higher magnification below. Yellow dashed lines in the magnified merge images indicate the paths used for the line-scan profiles shown in (E). Scale bars, 10 µm for full-cell images and 2 µm for magnified regions. **(E)** Representative line-scan intensity profiles showing the spatial distribution of endogenous KANK1 relative to mNG-Paxillin-positive FAs. Profiles were measured along the yellow dashed lines shown in (D). Fluorescence intensities were normalized independently for each channel to the maximum value within the corresponding line scan. In control cells, KANK1 peaks flank the paxillin-enriched adhesion core, consistent with peri-adhesion belt-like localization. Upon CCDC66 depletion, this spatial relationship is disrupted, with KANK1 signal appearing broader, displaced, or redistributed relative to paxillin-positive adhesions. **(F)** Representative confocal images of endogenous KANK1 (magenta) and phalloidin-stained F-actin (green) in U2OS cells transfected with control or CCDC66 siRNAs. Nuclei were counterstained with DAPI (blue). In control cells, KANK1 appears as discrete peripheral puncta near actin-bundle ends, whereas CCDC66-depleted cells show redistribution of KANK1 into elongated, fiber-like peripheral structures associated with bundled actin-rich regions. Upper panels show full-cell views, and lower panels show magnified peripheral regions. Scale bars, 10 µm and 2 µm, respectively. **(G)** Representative phase-contrast images of wound-healing assays in U2OS cells transfected with control, CCDC66, KANK1, or combined CCDC66 and KANK1 siRNAs. Confluent monolayers were scratched at 48 h post-transfection and imaged at 0 and 24 h post-wounding. Purple lines outline the migrating wound edges. Scale bars, 100 µm. **(H)** Quantification of wound closure from the assay shown in (G). Wound closure was calculated as the percentage reduction in wound area relative to the initial wound area at 0 h. Data are presented as mean ± S.E.M. from *n = 3* independent experiments. Symbols represent independent experiments. Fold changes and exact p values are indicated above the graph. Asterisks denote statistical significance (*p < 0.05, **p < 0.01, ns = not significant; two-way ANOVA with Tukey’s post hoc test).

Among these candidates, KANK1 was of particular interest because it binds talin at the FA rim and recruits cortical MT-stabilizing assemblies to promote MT capture near adhesions [24-27]. Moreover, KANK1 depletion phenocopies the enlarged FAs and impaired migration observed upon CCDC66 depletion [24, 26, 47]. Because KANK1 is not known to directly crosslink actin and MTs, the actin-MT crosslinking activity of CCDC66 suggested a possible mechanism by which KANK-dependent MT capture could be coupled to actin organization at adhesion sites. We therefore tested whether CCDC66 associates with KANK1 and contributes to its organization at FAs.

We first tested whether CCDC66 and KANK1 physically associate in cells. In pull-down experiments from cell lysates, FLAG-CCDC66 co-precipitated with endogenous KANK1 (Fig. 5B). We reciprocally confirmed this interaction using co-immunoprecipitation assays, which demonstrated that GFP-KANK1, but not the GFP control, co-precipitated FLAG-CCDC66 (Fig. 5C). We next mapped the KANK1 region required for CCDC66 association. KANK1 contains an N-terminal KN motif that mediates talin binding, a central coiled-coil region that binds liprin-β1, and a C-terminal ankyrin-repeat region that recruits KIF21A to cortical MT capture sites (Fig. S5A) [24, 25]. Using GFP-tagged KANK1 truncations, we found that FLAG-CCDC66 was recovered most strongly with the ankyrin-repeat fragment, whereas the coiled-coil, N-terminal, and KN fragments showed little or no enrichment above GFP background control (Fig. S5A,B). These data suggest that CCDC66 preferentially associates with the KANK1 region linked to MT-capture machinery.

We next asked whether CCDC66 affects KANK1 organization at peripheral adhesion sites. In U2OS cells expressing mNG-Paxillin, endogenous KANK1 accumulated adjacent to subsets of peripheral paxillin-positive FAs, forming the expected peri-adhesion belt-like pattern (Fig. 5D). Line-scan profiles confirmed that KANK1 flanked the paxillin-enriched adhesion core in control cells (Fig. 5E). Upon CCDC66 depletion, this spatial organization was disrupted. KANK1 no longer formed a defined peri-adhesion belt around paxillin-positive adhesions; instead, the KANK1 signal was displaced from the adhesion edge or redistributed into elongated peripheral structures (Fig. 5D,E). This redistribution was also evident relative to the actin cytoskeleton, where KANK1 shifted from discrete peripheral puncta near actin-bundle ends to fiber-like structures associated with bundled actin-rich regions (Fig. 5F). A similar disruption of the KANK1 belt was observed in cells expressing mNG-KANK1 (Fig. S5C,D). Together, these observations indicate that CCDC66 supports the normal peri-adhesion organization of KANK1 at peripheral FAs.

To investigate the functional relationship between CCDC66 and KANK1 during cell migration, we performed an epistasis analysis using a wound-healing assay, depleting each protein alone or in combination. First, we confirmed efficient KANK1 depletion by immunoblotting and qRT-PCR (Fig. S5E,F). Individual depletion of CCDC66 or KANK1 significantly impaired cell motility, reducing wound closure by ∼40% and ∼30% relative to control cells, respectively (Fig. 5G,H). Co-depletion of CCDC66 and KANK1 did not further reduce wound closure beyond either single-depletion condition (Fig. 5H). This non-additive phenotype indicates that CCDC66 and KANK1 operate within a shared functional pathway to support cell migration.

Collectively, these results support a model in which CCDC66 associates with KANK1 and helps maintain its peri-adhesion belt-like organization, thereby linking adhesion-associated MT capture to actin-MT crosstalk during cell migration.

### An unbiased transcriptomic screen identifies a CCDC66-associated KANK/ROCK cytoskeletal module linked to prognostic outcome

Having defined a CCDC66-dependent pathway that regulates adhesion remodeling and migration, we next asked whether this pathway is reflected in human tumors. To this end, we performed a transcriptome-wide expression-concordance analysis across 33 The Cancer Genome Atlas (TCGA) tumor cohorts, in which each gene was scored according to the strength of its correlation with CCDC66 expression within each cohort. Because our mechanistic data implicated cytoskeletal organization, MT regulation, adhesion remodeling, and migration, we compared a predefined set of GO categories related to these processes with all remaining GO terms. In representative cohorts, including chromophobe renal cell carcinoma (KICH) and sarcoma (SARC), these selected categories were shifted toward higher concordance scores than the background GO-term distribution, supporting focused analysis of adhesion- and migration-associated processes (Fig. S6A).

Within this focused set, “podocyte cell migration” emerged as a recurrently concordant term, ranking fifth in KICH and twentieth in SARC cohorts. Genes contributing to this term included several components of the pathway defined in U2OS cells, including CCDC66, KANK1, KANK2, ROCK1, and DAAM2 (Fig. S6B). These results provided an unbiased basis for defining a targeted CCDC66/KANK/ROCK-associated module for pan-cancer correlation and survival analyses.

Across the TCGA cohorts analyzed, CCDC66 showed positive associations with KANK1, KANK2, ROCK1, ROCK2, and DAAM2, indicating that this CCDC66/KANK/ROCK-associated module is transcriptionally coordinated in multiple tumor contexts (Fig. 6A, Table S1). Among these cohorts, KICH showed one of the strongest and most coherent correlation patterns, and we therefore prioritized it for outcome analysis. To determine whether clinical information was captured by CCDC66 alone or by the broader module, we first examined CCDC66 single-gene survival associations across the screened cohorts. This single-gene analysis showed that CCDC66 expression alone did not stratify overall survival in most screened cohorts, with adrenocortical carcinoma (ACC), a rare malignancy of the adrenal cortex, as the only significant exception (Table S2). Consistent with this pattern, CCDC66 expression alone did not significantly stratify overall survival in KICH (Fig. 6B; Table S2). By contrast, unsupervised grouping of KICH tumors based on the six-gene module identified two patient clusters with significantly different overall survival (Fig. 6C, D). The poorer-survival cluster showed coordinated higher expression of module components, indicating that prognostic information in KICH emerges from the integrated cytoskeletal module rather than from CCDC66 expression in isolation.

**Figure 6.**
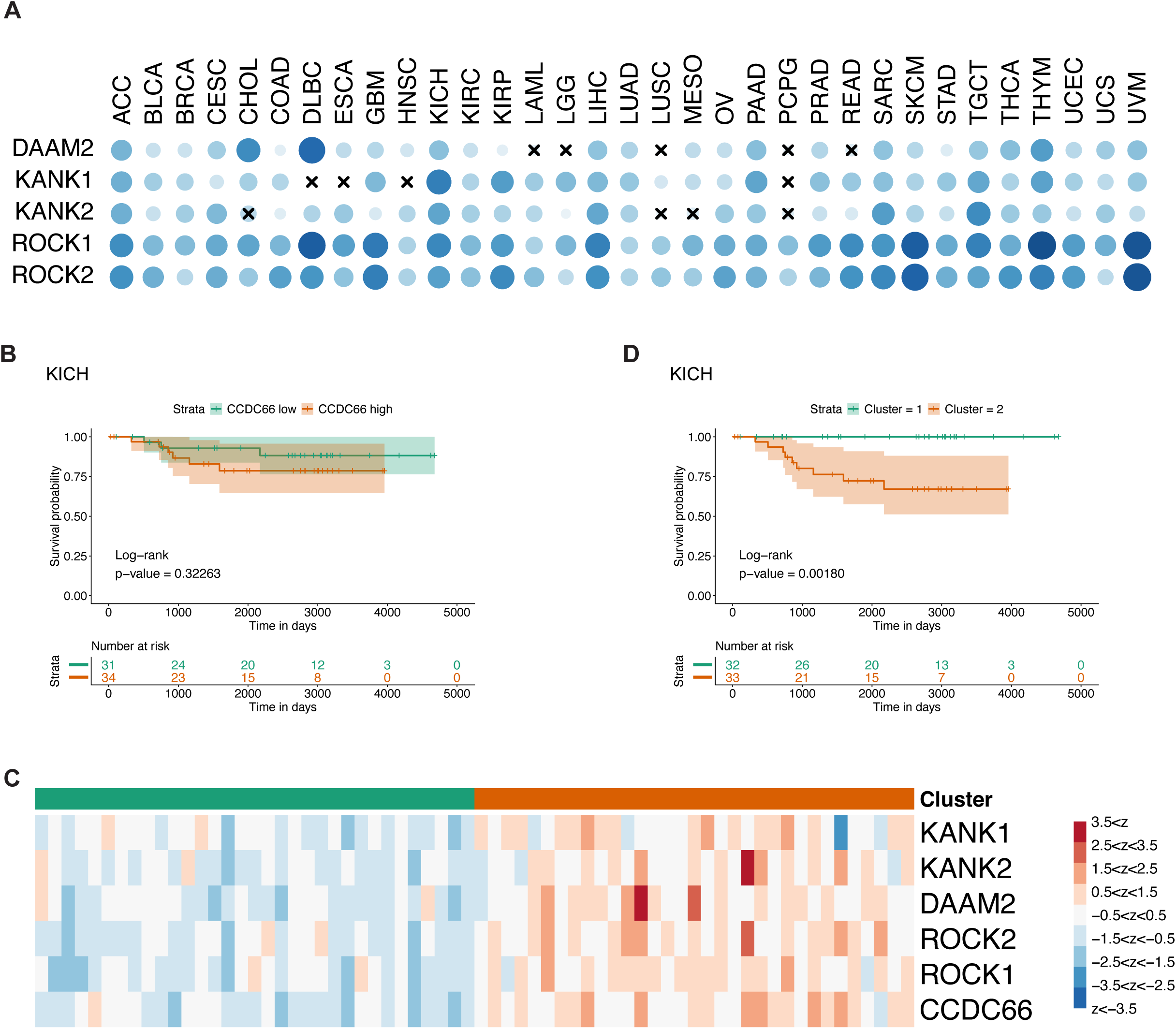
A CCDC66-associated KANK/ROCK cytoskeletal module is transcriptionally coordinated across tumors and linked to poor outcome in KICH. **(A)** Pan-cancer correlation analysis of CCDC66 with DAAM2, KANK1, KANK2, ROCK1, and ROCK2 across screened TCGA tumor cohorts. Blue circles indicate positive Spearman correlations, with circle size and color intensity reflecting correlation strength; crosses indicate non-significant associations. **(B)** Kaplan-Meier analysis of overall survival in kidney chromophobe carcinoma (KICH) patients stratified into CCDC66-low and CCDC66-high groups. CCDC66 expression alone did not significantly separate survival groups. The number of patients in each group is indicated in the plot. P value was calculated by log-rank test. **(C)** Heatmap of z-scored expression values for CCDC66, KANK1, KANK2, ROCK1, ROCK2, and DAAM2 in KICH tumors grouped by unsupervised clustering. Colors indicate normalized gene expression, with module-high and module-low clusters indicated by the top annotation bar. **(D)** Kaplan-Meier analysis of overall survival for the two KICH clusters shown in (C). The cluster enriched for the broader CCDC66/KANK/ROCK-associated cytoskeletal program showed significantly poorer overall survival, indicating that coordinated variation in the module, rather than CCDC66 expression alone, captures the prognostic behavior of this pathway. The number of patients in each group is indicated in the plot. P value was calculated by log-rank test.

Because our mechanistic studies were performed in U2OS osteosarcoma cells, we also examined the TCGA SARC cohort. CCDC66-high tumors showed a non-significant trend toward poorer overall survival (Fig. S6C, Table S2), and although the six-gene module separated SARC tumors into module-low and module-high expression groups (Fig. S6D), this grouping did not significantly stratify survival (Fig. S6E). Thus, the clinical association of the CCDC66/KANK/ROCK module is context dependent, with KICH providing the clearest evidence for outcome association.

Together, these transcriptomic analyses complement our cellular and biochemical data by showing that the CCDC66-regulated adhesion and cytoskeletal pathway is represented in human tumors as a coordinated CCDC66/KANK/ROCK-associated module. The KICH survival association aligns closely with our cellular findings, where CCDC66 depletion impaired motility, while its overexpression enhanced wound closure and Matrigel invasion. Integrating these clinical and cellular data, we propose a working model in which CCDC66 acts at adhesion-proximal actin-MT interfaces. Here, it links KANK-associated MT targeting to FA turnover and ROCK-driven contractility to coordinate cell migration (Fig. 7).

**Figure 7.**
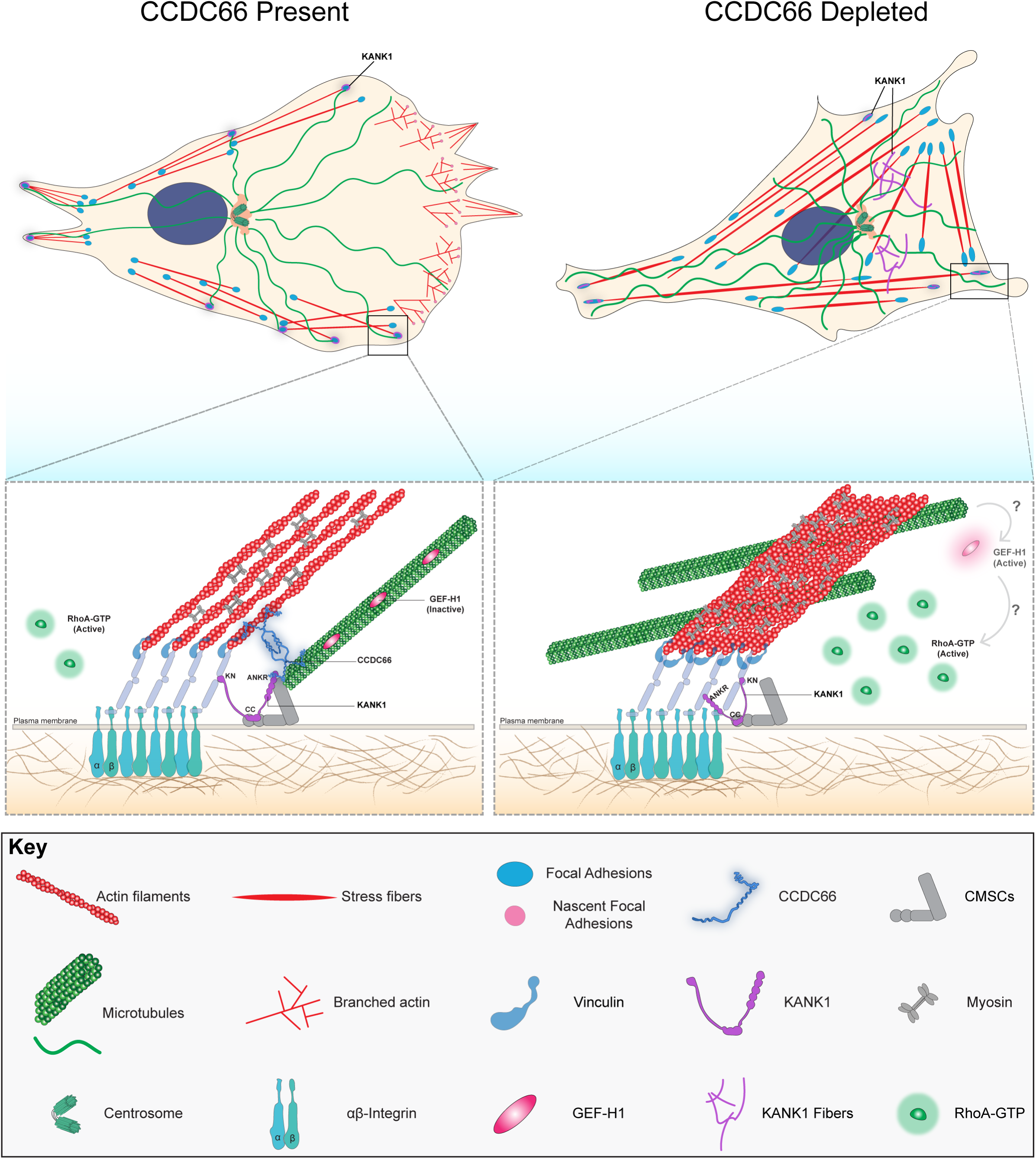
Working model for CCDC66 function at adhesion-proximal actin-microtubule interfaces. Model depicting how CCDC66 coordinates actin-MT coupling, KANK1-associated MT targeting, FA turnover, and RhoA-ROCK-dependent contractility during cell migration. In control cells, CCDC66 couples actin networks to microtubules at the focal adhesions. CCDC66 directly crosslinks actin filaments and MTs in vitro, associates with KANK1 in cells, and supports the peri-adhesion organization of KANK1 near peripheral FAs. This promotes efficient MT targeting to FAs, supports FA turnover, and helps maintain the balance between lamellipodial protrusion and actomyosin contractility. Upon CCDC66 depletion, MT targeting to FAs is reduced and KANK1 peri-adhesion organization is disrupted, with KANK1 redistributing into broader or fiber-like peripheral structures. Reduced adhesion-proximal MT targeting may weaken local MT-dependent restraint of RhoA signaling, potentially through altered GEF-H1 regulation. This promotes increased RhoA-ROCK activity, actomyosin contractility, stress fiber accumulation, and formation of enlarged, long-lived FAs. The resulting contractile state, marked by enlarged and long-lived FAs, limits Rac/Arp2/3-dependent protrusion and impairs efficient migration. Question marked arrows indicate proposed mechanisms that remain to be tested.

## Discussion

Efficient cell migration requires precise coordination of actin-driven protrusion, MT targeting, and FA turnover. In this study, we identified CCDC66 as a cytoskeletal organizer of this coordination. CCDC66 directly crosslinked actin and MTs *in vitro*, promoted MT targeting to FAs in cells, and associated with KANK1, positioning it as a link between actin-MT coupling and adhesion-associated MT targeting machinery. Functionally, CCDC66 supported FA turnover, sustained lamellipodial dynamics, and maintained the protrusion-contractility balance required for productive movement. These findings define a previously unrecognized CCDC66-dependent actin-MT-KANK1 axis and expand the non-ciliary functions of CCDC66 to cytoskeletal mechanisms relevant to adhesion remodeling and migration.

In migrating cells, Rac/Arp2/3-dependent branched actin networks drive lamellipodial protrusion, while RhoA/ROCK signaling and formin-dependent actin assembly promote stress fibers, actomyosin contractility, and adhesion maturation [3, 4, 9]. Our data indicate that CCDC66 helps maintain the balance between these opposing states. CCDC66 depletion shifted cells toward RhoA/ROCK-dependent contractile organization, marked by increased RhoA-GTP and p-MLC, bundled stress fibers, enlarged focal adhesions, and impaired lamellipodial dynamics. This state was accompanied by broadly slowed FA turnover: both assembly and disassembly rates were decreased, while FA lifetime increased. Reduced assembly is consistent with fewer productive leading-edge adhesion-forming events, whereas reduced disassembly explains why adhesions persist and remain coupled to bundled stress fibers long enough to enlarge. ROCK inhibition suppressed the contractile and adhesion phenotypes and restored efficient migration, and similarly, formin inhibition reduced excess linear actin and FA accumulation and improved wound closure. In contrast, constitutively active Rac1 failed to restore broad lamellipodial protrusions or efficient motility, indicating that the defect was not simply due to insufficient Rac1 activation. Together, these findings suggest that CCDC66 loss creates a ROCK-dependent contractile state with enlarged, long-lived FAs that prevents Rac/Arp2/3 signaling from generating sustained leading-edge protrusion.

These findings fit with current models in which MT capture at FAs coordinates adhesion remodeling with local RhoA regulation. Dynamic MTs target peripheral adhesions and promote FA turnover, whereas talin-KANK complexes at the FA rim connect adhesions to cortical MT-stabilizing assemblies containing liprins, ELKS, LL5β, CLASPs, and KIF21A [10-12]. MTs also restrain contractility through GEF-H1, a RhoA GEF that is inactive when bound to polymerized MTs and can activate RhoA-ROCK-myosin signaling when released during MT remodeling or depolymerization [14, 48-50]. Disruption of FA-associated MT capture, including KANK-dependent targeting, has been linked to GEF-H1 release, increased myosin activity, and actomyosin-driven FA growth and stabilization [50, 51]. In this context, reduced MT targeting after CCDC66 depletion provides a potential mechanism for increased RhoA-ROCK signaling and enlarged, long-lived adhesions. This model predicts that CCDC66 normally helps maintain adhesion-proximal MT targeting, thereby restraining GEF-H1-dependent RhoA activation and the resulting actomyosin-driven FA enlargement and persistence. Future work using GEF-H1 localization, activity reporters, or loss-of-function approaches will be important to test this mechanism directly.

Our results also refine current models of actin-MT coordination at adhesion sites. Previous studies identified mechanisms that guide or retain MT plus ends near FAs. EB proteins and spectraplakins such as MACF1/ACF7 guide growing MTs along actin structures toward adhesions, while KANK-dependent cortical complexes capture and stabilize MT plus ends near the FA rim [11, 19-21] [22, 24, 51]. These mechanisms explain how MTs are guided and positioned near adhesions, but not how FA-associated MT capture is physically integrated with force-bearing actin networks during adhesion remodeling. CCDC66 adds a distinct component to this interface by combining direct actin-MT crosslinking activity with KANK1 association and support of KANK1 organization at peripheral adhesion-associated sites. Although our functional validation focused on KANK1, KANK2 is also a characterized KANK-family regulator of adhesion-associated MT capture and was identified in both CCDC66 proximity interactomes and the CCDC66-associated tumor module [23, 30-32]. Future work should test whether CCDC66 also associates with KANK2 and whether KANK1 and KANK2 act redundantly or in context-specific ways during CCDC66-dependent MT targeting and FA remodeling.

Our transcriptomic analyses place the CCDC66-dependent cytoskeletal pathway in a broader disease context without reducing it to a single-gene marker. In KICH, a renal epithelial cancer, CCDC66 expression alone did not stratify survival, whereas coordinated expression of the CCDC66/KANK/ROCK-associated module did, with the poorer-survival group enriched for module components. This module-level association is consistent with our functional data showing that CCDC66 loss suppresses migration, whereas increased CCDC66 expression enhances wound closure and Matrigel invasion. In sarcoma (SARC), CCDC66-high tumors showed a non-significant trend toward poorer survival, but the grouped module did not stratify outcome. Thus, the clinical impact of this pathway is context dependent and may reflect differences in tumor lineage, tissue architecture, cohort heterogeneity, or the relative contribution of CCDC66 functions in migration, adhesion remodeling, and cell division. Future work should test whether module-high tumors show elevated ROCK activity, altered FA organization, increased invasion, or mitotic vulnerabilities in cancer models matched to the relevant tumor contexts.

More broadly, our findings expand the biology of CCDC66 beyond its established roles in cilia and mitosis. Because CCDC66 has been linked to retinal, olfactory, and Joubert neurodevelopmental phenotypes, its disease relevance may not be limited to canonical ciliary defects. Although we did not test disease models here, our data suggest that impaired actin-MT integration, FA turnover, and adhesion-dependent migration should be considered as potential non-ciliary contributors to CCDC66-associated pathology, particularly in contexts such as neuronal migration, polarization, and tissue organization.

In summary, our results identify CCDC66 as an actin-MT crosslinking factor that links KANK-associated MT targeting to FA turnover and efficient migration (Fig. 7). By supporting KANK1 organization at adhesion-proximal sites and limiting excessive ROCK-dependent FA enlargement and persistence, CCDC66 maintains the cytoskeletal balance required for productive protrusion and cell movement. This model expands the functional repertoire of CCDC66 beyond ciliary biology and provides a framework for investigating how disrupted actin-MT integration contributes to developmental disease and tumor-associated motility.

## Materials and Methods

### Plasmids and cloning

All plasmids used in this study are listed in Table S3. mScarlet was PCR amplified and cloned into pCDH-EF1-T2A-Puro using XbaI and EcoRI sites to generate pCDH-EF1-mScarlet-T2A-Puro. Human paxillin/PXN cDNA was PCR-amplified from pEGFP-Paxillin and cloned into pCDH-EF1-mNG-T2A-Puro or pCDH-EF1-mScarlet-T2A-Puro using EcoRI and NotI sites to generate mNG-Paxillin and mScarlet-Paxillin. Full-length human KANK1 was cloned into pCDH-EF1-mNG-T2A-Puro using EcoRI and NotI sites. For GFP-trap pulldown assays, full-length KANK1 and KANK1 fragments encoding the KN motif, N-terminal region, coiled-coil region, and ankyrin-repeat region were cloned into pEGFP-C1 using EcoRI and KpnI sites. mNG-RhoA WT and mNG-Rac1 Q61L were generated by PCR amplification of the corresponding coding sequences and cloned into pCDH-EF1-mNG-T2A-Puro using EcoRI and NotI sites. siRNA-resistant mNG-CCDC66 was generated by amplifying the siRNA-resistant CCDC66 coding sequence from GFP-CCDC66RR and cloning it into pCDH-EF1-mNG-T2A-Puro using NotI (from Batman, U., et al., 2022). For inducible expression, mNG and siRNA-resistant mNG-CCDC66 were PCR-amplified from the aforementioned pCDH-EF1-mNG-T2A-Puro constructs and introduced into the pDONR221 vector utilizing a BP recombination reaction. These entry clones were then recombined into the pInducer20 destination vector via an LR recombination reaction. N-FLAG-BirA*-CCDC66 was generated by Gateway LR recombination between pDONR221-CCDC66 (Conkar, D., 2017) and the N-FLAG-BirA* destination vector. All PCR-amplified inserts and cloning junctions were verified by Sanger sequencing before use.

### Cell culture, transfection, pharmacological treatments and lentiviral transduction

All cell lines, culture reagents, and plasmids used for transfection and lentiviral production are listed in Table S3. U2OS human osteosarcoma cells and HEK293T human embryonic kidney cells were cultured in DMEM supplemented with 10% fetal bovine serum and 1% penicillin-streptomycin. hTERT-RPE1 human retinal pigment epithelial cells were cultured in DMEM/F-12 supplemented with 10% fetal bovine serum and 1% penicillin-streptomycin. All cell lines were maintained at 37°C in a humidified incubator with 5% CO₂. Cell lines were routinely tested negative for mycoplasma contamination using a PCR-based assay. Where indicated, cells were treated with 10 µM Y-27632 (ROCK inhibitor), 10 µM SMIFH2 (Formin inhibitor), 40 µM CK-666 (Arp2/3 inhibitor), or 1 µg/ml Mitomycin C.

HEK293T cells were transfected with the indicated plasmids using 1 µg/µl 25-kDa polyethyleneimine (PEI). For transient expression and immunoprecipitation experiments, cells were processed 48 h after transfection. Recombinant lentiviruses were produced in HEK293T cells by co-transfecting the indicated lentiviral transfer vector with psPAX2 and pMD2.G packaging plasmids using PEI. Viral supernatants were collected 48 h after transfection, filtered through 0.45-µm filters, and titered on U2OS cells using a GFP-expressing control virus. To generate stable cell lines, U2OS cells were seeded in 6-well plates at 3 × 10⁵ cells per well, transduced at an estimated multiplicity of infection of 1 for 48 h, and selected with 1 µg/ml puromycin for 4 days before expansion and use in experiments.

### RNA Interference

All siRNAs used in this study are listed in Table S3. Cells were seeded at 30-40% confluency and transfected with 50 nM siRNA using Lipofectamine RNAiMAX in Opti-MEM according to the manufacturer’s instructions. Unless otherwise indicated, two sequential siRNA transfections were performed, and cells were analyzed 48 h after transfection or at the assay-specific endpoint. Knockdown efficiency was validated by qRT-PCR and, where indicated, by immunoblotting.

For double-depletion wound-healing assays, the total siRNA concentration was kept constant at 50 nM across all conditions. Cells were transfected with siControl + siControl, siControl + siCCDC66, siControl + siKANK1, or siCCDC66 + siKANK1, using equal amounts of each siRNA component. For rescue experiments, U2OS cells stably expressing mNG or siRNA-resistant mNG-CCDC66 (under stable or inducible promoters) were transfected with control or CCDC66 siRNAs using the same protocol and then processed for live-cell imaging or immunofluorescence. For inducible cell lines, doxycycline was added at a final concentration of 1 µg/mL 24 hours post-transfection and replenished every 24 hours until the completion of the experiment. For phenotypic quantification, cells with excessive fusion protein overexpression or abnormal subcellular localization were excluded.

### Immunofluorescence and confocal microscopy

Cells grown on coverslips were washed twice with PBS and fixed either with ice-cold methanol or with 4% PFA in cytoskeletal buffer containing 10 mM PIPES, 3 mM MgCl₂, 100 mM NaCl, 300 mM sucrose, 5 mM EGTA, and 0.1% Triton X-100, pH 6.9, for 10 min at room temperature. PFA fixation was used for paxillin, β-catenin, and phospho-myosin light chain staining, whereas methanol fixation was used for α-tubulin, γ-tubulin, and PCM1 staining. After fixation, coverslips were washed three times with PBS, permeabilized with 0.2% Triton X-100 in PBS for 5 min, and blocked for 1 h at room temperature in 3% BSA in PBS containing 0.1% Triton X-100. Primary antibodies were incubated for 2 h at room temperature or overnight at 4°C. Coverslips were then washed three times with PBS and incubated for 1 h at room temperature with Alexa Fluor-conjugated secondary antibodies, DAPI, and, where indicated, fluorescent phalloidin. After three final PBS washes, coverslips were mounted in Mowiol mounting medium containing N-propyl gallate. All primary antibodies, secondary antibodies, dyes, and working dilutions are listed in Table S3.

Confocal imaging was performed on a Leica Stellaris 8 inverted laser-scanning confocal microscope equipped with 405-, 488-, 561-, and 633-nm laser lines, one photomultiplier tube detector, three hybrid detectors, and an environmental chamber for live-cell imaging. Fixed-cell images were acquired using an HC PL APO CS2 63×/1.4 NA oil-immersion objective with the pinhole set to 1 Airy unit. Z-stacks of 10-20 optical sections were acquired with 0.2-0.5 µm spacing using 400-Hz bidirectional scanning and line averaging of 2-3. High-resolution images were acquired at 2096 × 2096 pixels with 1.5× optical zoom. Image acquisition was controlled using Leica Application Suite X software, and deconvolution was performed using the LAS X Lightning module where indicated. Within each experiment, images used for quantitative comparison were acquired with identical settings.

### Migration assays

#### Wound healing assay

U2OS cells were seeded into 4-well imaging chambers and transfected the next day with the indicated siRNAs. For rescue experiments, stable U2OS lines expressing mNG, siRNA-resistant mNG-CCDC66, or mNG-Rac1 Q61L were used. At 48 h post-transfection, confluent monolayers were scratched with a sterile 10-µl pipette tip, washed twice with PBS, and incubated in serum-free Opti-MEM. Where indicated, inhibitors or vehicle control were added immediately after scratching.

Wound healing was imaged on a Leica DMi8 inverted microscope equipped with an environmental chamber maintained at 37°C and 5% CO₂. Images were acquired every 2 h for 24 h using a 20× dry objective. At each time point, z-stacks of 5-10 optical sections were acquired with 1.86 µm spacing. All images were acquired in a 512 × 512 pixel format. Wound areas were measured in Fiji/ImageJ by manual tracing of the cell-free region. Wound closure was calculated as the percentage reduction in wound area at 24 h relative to the wound area at 0 h. For time-course analyses, wound area at each time point was normalized to the initial wound area at 0 h.

#### Random migration assay

U2OS cells were seeded in 24-well plates and transfected with control or CCDC66 siRNAs. At 48 h post-transfection, cells were trypsinized and re-seeded at low density, approximately 75,000 cells per well, into fibronectin-coated 2-well imaging chambers. Chambers were coated with fibronectin at 1 µg/cm² before cell seeding. After overnight attachment and spreading, live-cell imaging was performed using a Leica DMi8 inverted microscope equipped with an environmental chamber maintained at 37°C and 5% CO₂. Images were acquired every 6 min for 18 h using a 20× dry objective. At each time point, z-stacks of 5-10 optical sections were acquired with a 1.86-µm step size to maintain cells in focus during migration. Images were acquired at 512 × 512 pixels, and automated multiposition acquisition was controlled using LAS X software.

Cell trajectories were analyzed in Fiji/ImageJ using the Manual Tracking plugin. Individual cells were tracked frame by frame using the approximate cell center as the reference point. XY coordinate data were imported into the Ibidi Chemotaxis and Migration Tool to calculate accumulated distance, Euclidean distance, velocity, and directness. Migration trajectories were plotted in R/RStudio. Reagents, chamber details, and coating materials are listed in Table S3.

#### Transwell migration and invasion assays

For migration assays, U2OS cells were seeded in 6-well plates at approximately 40% confluency and transfected the following day with control or CCDC66 siRNAs. At 48 h post-transfection, cells were trypsinized, resuspended in serum-free Opti-MEM, and 3 × 10⁵ cells were added to each 8-µm pore-size Transwell insert. Complete growth medium containing 10% FBS was added to the lower chamber as a chemoattractant. Cells were allowed to migrate for 48 h at 37°C and 5% CO₂. For invasion assays, the upper surface of each Transwell insert was coated with 40 µl of Matrigel diluted 1:1 in serum-free DMEM and incubated for 2 h at 37°C to allow gelation. Prior to the assay, doxycycline-inducible U2OS cells expressing mNG or mNG-CCDC66 were cultured in 6-well plates and pre-treated with doxycycline (1 µg/mL) for 24 h to ensure construct expression at the time of seeding. These pre-induced cells were trypsinized, resuspended in serum-free Opti-MEM containing doxycycline, and 5 × 10⁵ cells were added to each Matrigel-coated insert. Complete growth medium containing 10% FBS and doxycycline was added to the lower chamber, and cells were allowed to invade for 96 h at 37°C and 5% CO_2_, with doxycycline replenished every 24 h throughout the duration of the experiment.

After migration or invasion, inserts were washed twice with PBS and fixed with 4% PFA. Cells remaining on the upper surface of the membrane were removed with a cotton swab. Membranes were excised, stained with DAPI, mounted in Mowiol, and imaged using a Leica DMi8 inverted fluorescence microscope with a 40×/1.3 NA oil objective. Z-stacks of 5-30 optical sections were acquired with a 0.49-µm step size at 1024 × 1024 pixels. For quantification, 10 random fields of view were acquired per membrane, and nuclei on the lower membrane surface were counted in Fiji/ImageJ. Results were expressed as the number of migrated or invaded cells per field. Reagent and insert details are listed in Table S3.

### Live-cell super-resolution imaging and analysis of cytoskeletal dynamics

#### Live-cell super-resolution imaging of focal adhesion dynamics

U2OS cells stably expressing mNG-Paxillin were transfected with control or CCDC66 siRNAs. At 48 h post-transfection, cells were trypsinized and re-seeded at low density (15% confluency) into 5 µg/cm² fibronectin-coated 2-well imaging chambers. After overnight spreading, focal adhesion dynamics were imaged on a Zeiss Elyra 7 Lattice SIM2 system maintained at 37°C and 5% CO_2_. Time-lapse images were acquired every 1 min for 1 h using a Plan-Apochromat 63×/1.4 NA Oil DIC M27 objective, 405, 488, 561, and 633 nm laser lines, standard excitation/emission filter sets and an incubation chamber for live-cell experiments and an sCMOS camera (version 4.2 CL HS). Raw images were reconstructed using the SIM module in ZEN Black software, and channel alignment was calibrated using TetraSpeck beads.

To quantify focal adhesion assembly and disassembly kinetics, SIM-reconstructed time-lapse images of mNG-Paxillin were first segmented in Aivia. Because of background fluorescence, a supervised pixel-classifier model was trained on representative frames to distinguish focal adhesion signal from background and generate a segmented focal adhesion channel. The segmented image sequences were imported into Fiji/ImageJ, where automated thresholding was applied to remove residual background. Focal adhesions were then tracked over time using the TrackMate plugin. Detection was performed with the thresholding detector using a threshold value of 100 and the “simplify contours” option enabled. Focal adhesion objects were linked between consecutive frames using the overlap tracker with a minimum intersection-over-union value of 0.1 and a scale factor of 1.

TrackMate output files containing focal adhesion area and temporal information were exported as CSV files and analyzed in R. Tracks persisting for fewer than seven consecutive frames were excluded to remove transient segmentation noise. For each retained track, the focal adhesion area profile was used to identify the maximum area. The assembly rate was calculated as the linear regression slope of focal adhesion area over time from initial appearance to maximum area. The disassembly rate was calculated as the slope from maximum area to the subsequent minimum area during the disassembly phase. Tracks without a clearly defined assembly phase or disassembly phase were assigned as not applicable and excluded from the corresponding rate calculation. FA lifetime was defined as the duration for which a tracked adhesion persisted across consecutive frames.

#### Live-cell imaging and analysis of actin protrusion dynamics

U2OS cells stably expressing RFP-LifeAct were seeded into 2-well imaging chambers at approximately 20% confluency and transfected with control or CCDC66 siRNAs. At 48 h post-transfection, cells were imaged on a Leica Stellaris 8 inverted confocal microscope maintained at 37°C and 5% CO₂. Single-plane time-lapse images were acquired every 10 s for 10 min using an HC PL APO CS2 63×/1.4 NA oil-immersion objective with the pinhole set to 1 Airy unit. Images were acquired at 1280 × 1280 pixels using a 400-Hz bidirectional scan mode. Image acquisition and real-time deconvolution were performed with LAS X software using the Lightning module.

Protrusion dynamics were quantified from kymographs generated in Fiji/ImageJ. Line ROIs were drawn manually perpendicular to the cell edge at distinct protrusive regions, and kymographs were generated using the Kymograph Builder plugin. Individual protrusion events were manually traced from the start to the end of the protrusive growth phase. Protrusion distance and persistence were calculated from the spatial and temporal components of each trace, respectively, using the calibrated image scale and acquisition interval. Protrusion velocity was calculated as protrusion distance divided by persistence. For each condition, 15-30 protrusion events from 7-10 cells were analyzed across three independent experiments.

#### Live-cell imaging and analysis of EB3 targeting to focal adhesions

U2OS cells stably expressing GFP-EB3 and mScarlet-Paxillin were seeded into 2-well imaging chambers at approximately 20% confluency and transfected with control or CCDC66 siRNAs. At 48 h post-transfection, cells were imaged on the same Leica Stellaris 8 confocal system under identical environmental conditions. Single-plane time-lapse images were acquired every 1 s for 1 min using an HC PL APO CS2 63×/1.4 NA oil-immersion objective with the pinhole set to 1 Airy unit. Images were acquired at 992 × 992 pixels using a 400-Hz bidirectional scan mode, and real-time deconvolution was performed with the LAS X Lightning module.

Microtubule plus-end targeting to FAs was quantified using Fiji/ImageJ and custom R scripts. Peripheral FAs were manually outlined in the mScarlet-Paxillin channel, and their XY coordinates, bounding boxes, and areas were exported. GFP-EB3 comets were tracked in the corresponding time-lapse sequences using TrackMate. Comets were detected with the Laplacian of Gaussian detector, with the detection threshold adjusted empirically for each cell to identify EB3 comets while excluding background. Tracks were reconstructed using the Linear Assignment Problem tracker with a maximum frame-to-frame linking distance of 0.8 µm. Gap closing was enabled with a maximum distance of 1.2 µm and a maximum gap of 2 frames.

EB3 targeting events were calculated by integrating the FA ROI data with TrackMate spot and track coordinates in R. A targeting event was defined as the intersection of a GFP-EB3 track with a defined peripheral FA region. Multiple intersections of the same EB3 track with the same FA were counted as a single targeting event using the unique TrackMate track ID. EB3 targeting rate was calculated as the number of unique EB3 targeting events normalized to FA area. FAs that received no EB3 targeting event during the 1-min acquisition were classified as non-targeted. Data were visualized in R using ggplot2 and ggforce, with individual measurements shown as Sina points overlaid on violin plots.

### Fixed cell image processing and analysis

#### Focal adhesion size and density

Cells were grown on coverslips, transfected with control or CCDC66 siRNAs, and fixed 48 h post-transfection as described above. For rescue experiments, U2OS cells stably expressing mNG or siRNA-resistant mNG-CCDC66 were processed in parallel. For pharmacological perturbation experiments, cells were treated with DMSO, Y-27632, or SMIFH2 before fixation. Focal adhesions were visualized either by endogenous paxillin immunostaining or by mNG-Paxillin fluorescence. When mNG-Paxillin was used, cells were stained for β-catenin to define cell boundaries.

Focal adhesion segmentation was performed using Aivia followed by Fiji/ImageJ. A supervised pixel-classifier model was trained in Aivia on representative images to distinguish focal adhesion signal from background. The segmented focal adhesion channel was imported into Fiji/ImageJ, and automated thresholding was applied to remove residual background. Focal adhesion area was measured using the Analyze Particles function with a minimum size threshold of 0.15 µm². Focal adhesion density was calculated as the number of focal adhesions per cell divided by total cell area. Cell area was measured by manually outlining the cell boundary using the Freehand selection tool in Fiji/ImageJ.

#### Stress fiber number and thickness

U2OS cells or U2OS cells stably expressing mNG-Paxillin were transfected with control or CCDC66 siRNAs, fixed 48 h post-transfection, and stained with fluorescent phalloidin to visualize F-actin. For inhibitor experiments, cells were treated with DMSO, Y-27632, or SMIFH2 before fixation. Stress fiber analysis was performed in Fiji/ImageJ. The phalloidin channel was thresholded to enhance filamentous actin structures. To quantify stress fiber number per cell, multiple freehand line profiles were drawn across the cell to intersect visible stress fibers in different orientations. Intensity profiles were generated for each line, and stress fiber peaks were counted using the Find Peaks plugin. Peak counts from all line profiles within a cell were summed to obtain the total stress fiber number per cell. To quantify stress fiber thickness, straight line profiles were drawn perpendicular to individual stress fibers, and the width of each corresponding intensity peak was measured using the calibrated image scale.

#### Microtubule targeting to focal adhesions

U2OS cells stably expressing mNG-Paxillin were transfected with control or CCDC66 siRNAs and fixed 48 h post-transfection. A sequential fixation protocol was used to preserve the mNG-Paxillin signal while maintaining α-tubulin staining quality. Cells were first fixed with ice-cold methanol for 3 min, washed three times with PBS, and then fixed with 4% PFA. Subsequent immunostaining for α-tubulin and DAPI was performed as described above. Analysis was restricted to peripheral focal adhesions located at the cell edge. A focal adhesion was scored as MT-targeted when a free MT end overlapped with the adhesion or was positioned in close proximity and oriented toward it. MT targeting efficiency was calculated for each cell as the percentage of targeted peripheral focal adhesions relative to the total number of peripheral focal adhesions analyzed.

#### Centrosome polarization

U2OS cells transfected with control or CCDC66 siRNAs were grown on coverslips and wounded 48 h post-transfection using a sterile 10-µl pipette tip. After 4 h of migration toward the wound edge, cells were fixed and stained for γ-tubulin and DAPI. Centrosome orientation was quantified in Fiji/ImageJ by measuring the angle between the nucleus-to-centrosome axis and a line perpendicular to the wound edge. The angle vertex was placed at the center of the nucleus, one arm was drawn toward the centroid of the γ-tubulin-positive centrosome, and the second arm was drawn perpendicular to the wound edge. Cells with centrosome angles between 0° and 90° were classified as polarized toward the wound, whereas cells with angles between 90° and 180° were classified as unoriented.

### *In vitro* reconstitution assays and TIRF imaging

*In vitro* actin-microtubule reconstitution assays were performed in 7-8 µl flow chambers assembled from silanized coverslips, double-sided tape, and glass slides. Coverslips were prepared as described previously with minor modifications [52]. Briefly, coverslips were incubated overnight in 12% hydrochloric acid at 60°C instead of piranha solution treatment, washed extensively, incubated in 0.1 M potassium hydroxide, dried, silanized with 0.05% dichlorodimethylsilane in trichloroethylene, washed, sonicated in methanol, rinsed with ultrapure water, and dried before chamber assembly.

Alexa Fluor 647-labeled stabilized microtubule seeds were prepared by mixing in-house purified bovine brain tubulin with in-house labeled tubulin at a final tubulin concentration of 1 mg/ml and 15% labeling in BRB80 buffer containing 1 mM GMP-CPP. BRB80 consisted of 80 mM K-PIPES, 1 mM MgCl_2_, and 1 mM EGTA, pH 6.8. The tubulin mixture was incubated on ice for 15 min, centrifuged at 90,000 rpm for 5 min at 4°C, snap-frozen, and stored at −80°C in 5 µl aliquots. On the day of imaging, aliquots were supplemented with BRB80 containing 1 mM GMP-CPP, polymerized for 1 h at 37°C, centrifuged at 13,000 rpm for 5 min, resuspended in BRB80 containing 1 µM Taxol, and incubated for an additional 1 h at 37°C.

Rhodamine-labeled rabbit muscle actin was polymerized according to the manufacturer’s instructions. Briefly, frozen actin stock was diluted to 1 mg/ml in General Actin Buffer containing 20 mM Tris-HCl, pH 8.0, 0.2 mM CaCl_2_, 0.2 mM ATP, and 1 mM DTT, and incubated on ice for 1 h. Polymerization was initiated by addition of one-tenth volume of 10× polymerization buffer containing 1 M KCl, 20 mM MgCl_2_, and 10 mM ATP, followed by incubation for 1 h at room temperature. Polymerized F-actin was diluted tenfold in 1× polymerization buffer before binding reactions.

For microtubule immobilization, anti-β-tubulin antibody diluted 1:10 in BRB80 containing 1 µM Taxol was introduced into the flow chamber. The surface was blocked with 5% Pluronic F-127 for 5 min and washed with at least five chamber volumes of BRB80 containing 2 mg/ml casein and 1 µM Taxol. Stabilized microtubule seeds were diluted 50-fold in BRB80 containing 1 µM Taxol, flowed into the chamber, and allowed to bind to the antibody-coated surface. Chambers were then further blocked with BRB80 containing 0.5 mg/ml casein.

For microtubule-binding reactions, 100 nM His-MBP-mNeonGreen or His-MBP-mNeonGreen-CCDC66 was introduced into chambers containing immobilized microtubules in BRB80 supplemented with 0.5 mg/ml casein and an oxygen-scavenging system consisting of 0.2 mg/ml glucose oxidase, 0.035 mg/ml catalase, 4.5 mg/ml glucose, and 140 mM β-mercaptoethanol. For actin-binding and actin-microtubule crosslinking reactions, rhodamine-labeled F-actin was pre-incubated with 100 nM His-MBP-mNeonGreen or His-MBP-mNeonGreen-CCDC66 for 5 min at room temperature in BRB80 containing 1 µM Taxol, 0.5 mg/ml casein, 0.1% methylcellulose, and the oxygen-scavenging system. The protein-actin mixture was then introduced into chambers with or without immobilized Alexa Fluor 647-labeled microtubules. Chambers were sealed with nail polish before imaging.

TIRF imaging was performed at room temperature on a Zeiss Elyra 7 Lattice SIM microscope operated in TIRF mode, equipped with a 100×/1.46 NA objective, LBF 405/488/561/642 filter set, and an ORCA-Fusion camera. Images were acquired using ZEN Black software with a 1024 × 1024 pixel field of view.

### RhoA activity assay

Active GTP-bound RhoA was quantified using a RhoA Activation Assay Biochem Kit. U2OS cells stably expressing mNG-RhoA were transfected with control or CCDC66 siRNAs in 6-cm dishes. At 48 h post-transfection, cells were washed with ice-cold TBS and lysed on ice in the kit-provided lysis buffer supplemented with protease inhibitors. Lysates were rapidly cleared by centrifugation at 10,000 × g for 1 min at 4°C to preserve GTP-bound RhoA. Protein concentrations were determined by Bradford assay, and lysates were normalized to equal total protein concentration and volume. An aliquot of each normalized lysate was retained as the total RhoA input. The remaining lysate was incubated with Rhotekin-RBD beads for 1 h at 4°C with rotation. Beads were washed three times with the kit-provided wash buffer, and bound proteins were eluted by boiling in SDS sample buffer. Input and RBD-bound samples were analyzed by immunoblotting. RhoA activity was quantified in Fiji/ImageJ by subtracting local background from each band. The RBD-bound RhoA signal was normalized to the corresponding total RhoA input, which was first normalized to β-actin. Values were expressed as fold change relative to siControl.

### Immunoprecipitation

Protein interaction experiments were performed in HEK293T cells transfected with the indicated plasmids. At 48 h post-transfection, cells were washed with PBS and lysed in LAP200 buffer containing 50 mM HEPES pH 7.4, 100 mM KCl, 1 mM EGTA, 1 mM MgCl_2_, 10% glycerol, and 0.3% NP-40, freshly supplemented with 1 mM PMSF and protease inhibitors. Lysates were incubated for 15 min at 4°C and cleared by centrifugation at 15,000 × g for 10 min at 4°C. An aliquot of cleared lysate was retained as input. The remaining lysate was incubated overnight at 4°C with rotation using either GFP-Trap agarose beads or anti-FLAG M2 affinity gel, as indicated for each experiment. Beads were washed three times with LAP200 buffer, and bound proteins were eluted in SDS sample buffer and analyzed by immunoblotting. For KANK1 domain-mapping experiments, relative pulldown efficiency was calculated from immunoblot band intensities as the amount of co-precipitated FLAG-BirA*-CCDC66 normalized to the corresponding FLAG-BirA*-CCDC66 input and recovered GFP-tagged KANK1 bait. Values were expressed relative to the GFP control where indicated.

### Cell lysis and immunoblotting

For whole-cell extracts, cells were washed with cold PBS and lysed in RIPA buffer supplemented with protease inhibitors for 30 min at 4°C. Lysates were cleared by centrifugation at 15,000 × g for 15 min at 4°C, and protein concentrations were determined by Bradford assay. Equal amounts of protein were resolved by SDS-PAGE and transferred to nitrocellulose membranes. Membranes were blocked for 1 h at room temperature in 5% non-fat dry milk in TBS-T and incubated overnight at 4°C with primary antibodies diluted in 5% BSA in TBS-T. After three washes in TBS-T, membranes were incubated for 1 h at room temperature with IRDye 680- or IRDye 800-conjugated secondary antibodies. Blots were washed three additional times and imaged using a LI-COR Odyssey infrared imaging system. Band intensities were quantified using Fiji/ImageJ with local background subtraction. Primary and secondary antibodies, including catalog numbers and working dilutions, are listed in Table S3.

### RNA isolation, cDNA synthesis and qRT-PCR

U2OS cells were seeded in 6-well plates and transfected with control, CCDC66, or KANK1 siRNAs. At 48 h post-transfection, total RNA was isolated using the NucleoSpin RNA kit according to the manufacturer’s instructions. RNA concentration and purity were assessed by absorbance measurements at 260 and 280 nm. cDNA was synthesized from 1 µg total RNA using the iScript cDNA synthesis kit. qRT-PCR was performed using GoTaq qPCR Master Mix and gene-specific primers listed in Table S3. Relative transcript levels were calculated using the ΔΔCt method, normalized to GAPDH, and expressed relative to the corresponding control siRNA condition.

### Mass spectrometry data analysis

Published CCDC66 proximity-labeling datasets were reanalyzed to identify candidate CCDC66-associated proteins [46, 53]. Protein fold-change values were log_2_-transformed, and proteins with log_2_(fold change) ≥ 1 were retained for downstream analysis. Candidate interactors were filtered using the CRAPome database, excluding proteins with a contamination frequency >50%. This threshold was selected to remove frequent contaminants while retaining known CCDC66-associated proteins. The remaining proteins were subjected to Gene Ontology (GO) enrichment analysis using DAVID, with enriched terms considered significant at P < 0.05. Based on the enriched GO categories and literature-supported annotation, proteins were assigned to four functional groups: cell migration, actin cytoskeleton organization, cell adhesion, and small GTPase-mediated signal transduction. Protein-protein interaction edges among the filtered candidate interactors were obtained from the STRING database, and the resulting interaction networks were visualized in Cytoscape.

### TCGA transcriptomic and survival analyses

Gene expression profiles of tumors were preprocessed by the unified RNA-Seq pipeline of TCGA consortium (https://portal.gdc.cancer.gov). For each cancer type, HTSeq-FPKM files of all primary tumors from the most recent data freeze (i.e., Data Release 45 - December 04, 2025) were downloaded. Metastatic tumors were not included since their underlying biology would be very different from primary tumors. The gene expression profiles of primary tumors were first log2-transformed and z-normalized within each cohort before further analysis. Clinical annotation files of cancer patients were used to extract their survival characteristics (i.e., days to last follow-up for alive patients and days to death for dead patients). For each cancer type, Clinical Supplement files of all patients from the most recent data freeze were downloaded.

To perform concordance analysis between a specific gene (CCDC66 in our case) and GO categories, GO biological processes, which includes 7538 gene sets, were downloaded (https://www.gsea-msigdb.org). The concordance score between a target gene and a GO category is defined as the geometric mean of absolute values of Spearman correlations between the target gene and genes in the GO category. A higher concordance score means the genes in the corresponding GO category are more concordantly expressed with the target gene. A predefined set of migration-, adhesion-, cytoskeleton-, centrosome-, cilium- and MT-associated GO categories (209 categories) was compared with all remaining GO categories (7329 categories) using the Mann-Whitney test.

To perform survival analysis using gene expression profiles, only patients with available survival information and gene expression profile were included. The heat maps of gene expression values were based on the z-normalized gene expression values. For computational analyses based on a gene module, samples were grouped into two categories using k-means clustering (k = 2) on the z-normalized gene expression values. Kaplan-Meier survival curves of these two groups were then compared. The P value obtained from the log-rank test performed on these two survival curves was displayed.

### Statistical analyses

Statistical analyses were performed using GraphPad Prism and R/RStudio, as indicated. Data are presented as mean ± S.E.M. unless otherwise stated. For cell-based experiments, n denotes independent biological experiments, as specified in each figure legend. When multiple individual measurements were collected within a single experiment (e.g., individual cells, focal adhesions, protrusion events, or EB3 tracks), these values were plotted to show the data distribution. For statistical testing, individual measurements were first averaged within each independent experiment, and tests were performed on these experiment-level means.

Two-condition comparisons were analyzed using paired or unpaired Student’s t tests, as specified in the figure legends. Experiments comprising multiple conditions (e.g. siRNA depletions, rescue lines, and inhibitor treatments) were analyzed using one-way or two-way analysis of variance (ANOVA) followed by Tukey’s multiple-comparisons test. Gene Ontology (GO) enrichment score distributions were compared using the Mann-Whitney U test. Survival differences were assessed using the log-rank test, and gene-expression correlations were calculated using Spearman’s rank correlation. Exact P values are reported in the figures where applicable. Statistical significance is indicated as ns (not significant) for P ≥ 0.05, or explicitly as P < 0.05, P < 0.01, P < 0.001, and P < 0.0001.

## Acknowledgements

We thank the members of CytoLab for their insightful feedback on this work. This project has received funding from the European Union’s Horizon Europe program under the European Research Council (ERC) Starting Grant agreement No. 101078097 (SatelliteHomeostasis) to ENF. This work was additionally supported by EMBO Installation Grant 3622, an EMBO Young Investigator Award, TÜBİTAK BİDEB grant 120C148, and TÜSEB grant 38629 to ENF.

## Author contributions

F.B.T. and E.N.F.-K. conceived the study and designed the experimental strategy. F.B.T. performed most experiments and analyses, including cell migration assays, siRNA depletion and rescue experiments, immunofluorescence, live-cell imaging, focal adhesion dynamics, cytoskeletal quantifications, and figure preparation. I.S. performed the pulldown experiments and Matrigel invasion assays together with F.B.T. J.D. performed the *in vitro* actin-microtubule reconstitution assays and TIRF imaging together with F.B.T. M.G. performed the TCGA transcriptomic, survival, and module-level analyses. E.N.F.-K. supervised the project, secured funding, and guided data interpretation. F.B.T. and E.N.F.-K. wrote the manuscript with input from all authors. All authors reviewed and approved the final manuscript.

---

**Figure S1.**
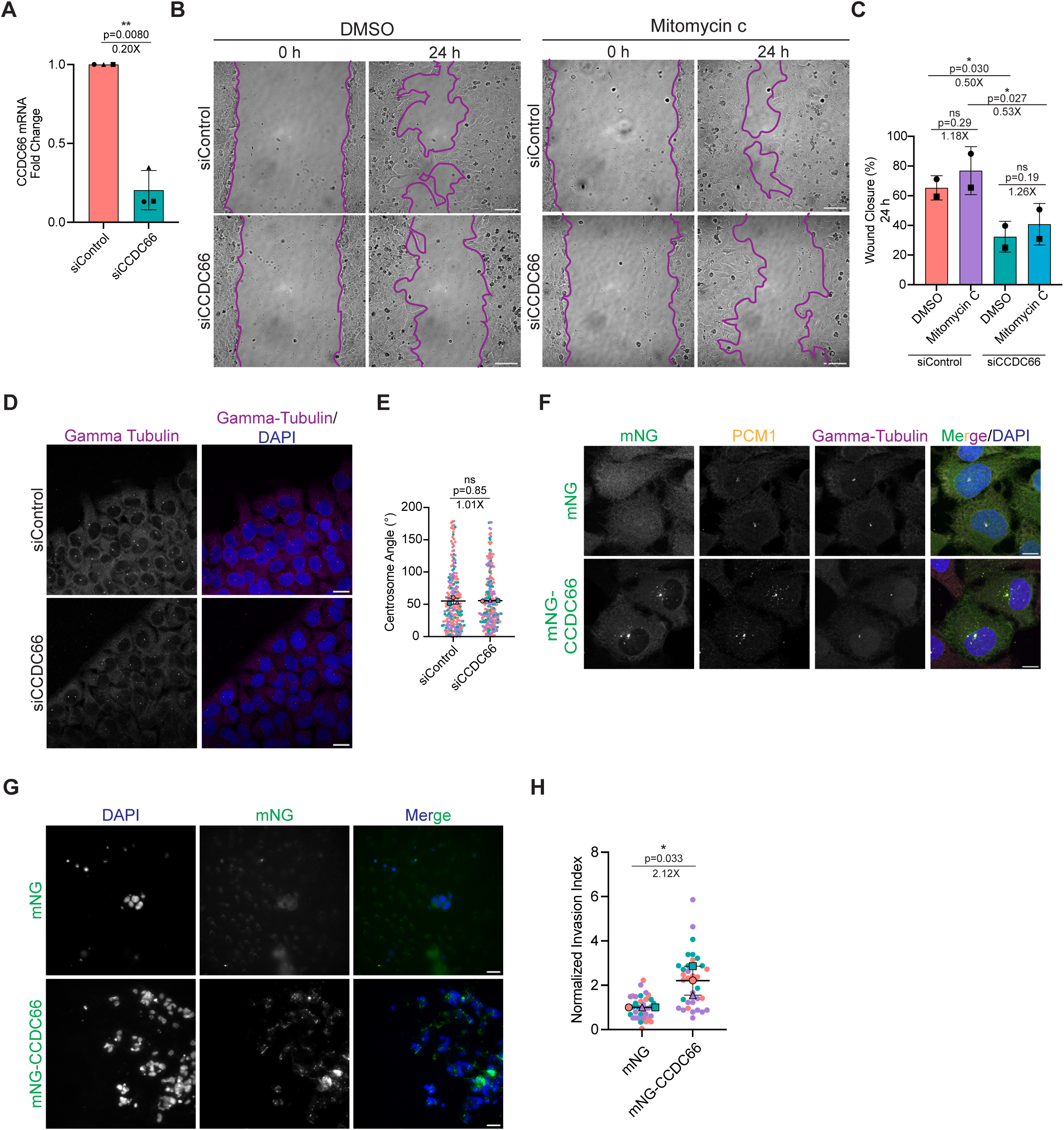
CCDC66 depletion and rescue construct validation, invasion analysis, and controls for proliferation and polarity. **(A)** Validation of CCDC66 knockdown efficiency at the mRNA level. U2OS cells were transfected with control or CCDC66 siRNAs. At 48 hours post-transfection, total RNA was extracted, reverse transcribed into cDNA, and analyzed by qPCR. The bar graph presents the fold change in CCDC66 mRNA expression relative to the control. Data are presented as mean ± S.E.M. from *n = 3* independent experiments. Dots represent independent experiments. Fold change relative to siControl and exact p values are indicated above the graph. Asterisks denote statistical significance (ns: not significant; paired Student’s *t* test). **(B)** Representative phase-contrast images of a wound healing assay in U2OS cells transfected with control or CCDC66-targeting siRNAs at 0 and 24 hours. At 48 hours post-transfection with control or CCDC66 siRNAs, cells were treated with either DMSO or 1 µg/ml Mitomycin C for 1 hour and washed. Confluent cell monolayers were then scratched, and cells were monitored via live-cell imaging every 2 hours over a 24-hour period. Purple lines outline the migrating wound edges. Scale bars, 100 µm. **(C)** Quantification of wound closure from the assay shown in (B). Wound closure was calculated as the percentage reduction in wound area relative to the initial wound area at 0 h. Data are presented as mean ± S.E.M. from *n = 2* independent experiments. Dots represent independent experiments. Fold changes and exact p-values are indicated above the graph. Asterisks denote statistical significance (*p < 0.05, ns: not significant; paired Student’s *t* test). **(D)** Representative immunofluorescence images illustrating centrosome polarization in migrating U2OS cells. At 48 hours post-transfection with control or CCDC66 siRNAs, confluent cell monolayers were wounded and incubated for 4 hours to allow for cellular reorientation towards the wound edge. Cells were then fixed and immunostained for the centrosome marker γ-Tubulin (magenta). Nuclei were counterstained with DAPI (blue). Scale bars, 20 µm. **(E)** Quantification of the centrosome positioning angle relative to the wound edge from the assay shown in (D). The centrosome positioning angle was measured between the nucleus-to-centrosome axis and the vector perpendicular to the wound edge. Cells with angles between 0° and 90° were scored as polarized toward the wound. Data are presented as mean ± S.E.M. from *n = 3* independent experiments. Colored dots represent individual measurements, with each color corresponding to one independent experiment Fold change relative to siControl and exact p values are indicated above the graph. Asterisks denote statistical significance (ns: not significant; paired Student’s *t* test). **(F)** Validation of doxycycline-inducible U2OS rescue lines expressing mNG or siRNA-resistant mNG-CCDC66. Cells were stained for PCM1 (yellow), gamma-tubulin (magenta), and DAPI (blue). mNG showed diffuse localization, whereas mNG-CCDC66 localized to centrosomes and PCM1-positive centriolar satellites. Scale bars, 10 µm. **(G)** Representative images from Matrigel-coated Transwell invasion assays using U2OS cells expressing mNG or mNG-CCDC66. Invaded cells on the lower surface of the membrane were fixed and stained with DAPI to visualize nuclei. mNG fluorescence confirms expression of the indicated constructs. Scale bars, 30 µm. **(H)** Quantification of Matrigel invasion from the assay shown in (G). Invasion was quantified as the number of DAPI-positive cells on the lower surface of the membrane and normalized to the mNG control condition. Data are presented as mean ± S.E.M. from *n = 3* independent experiments. Colored dots represent individual measurements, with each color corresponding to one independent experiment. Fold change relative to mNG and exact p value are indicated above the graph. Asterisks denote statistical significance (*p < 0.05; paired Student’s *t* test).

**Figure S2.**
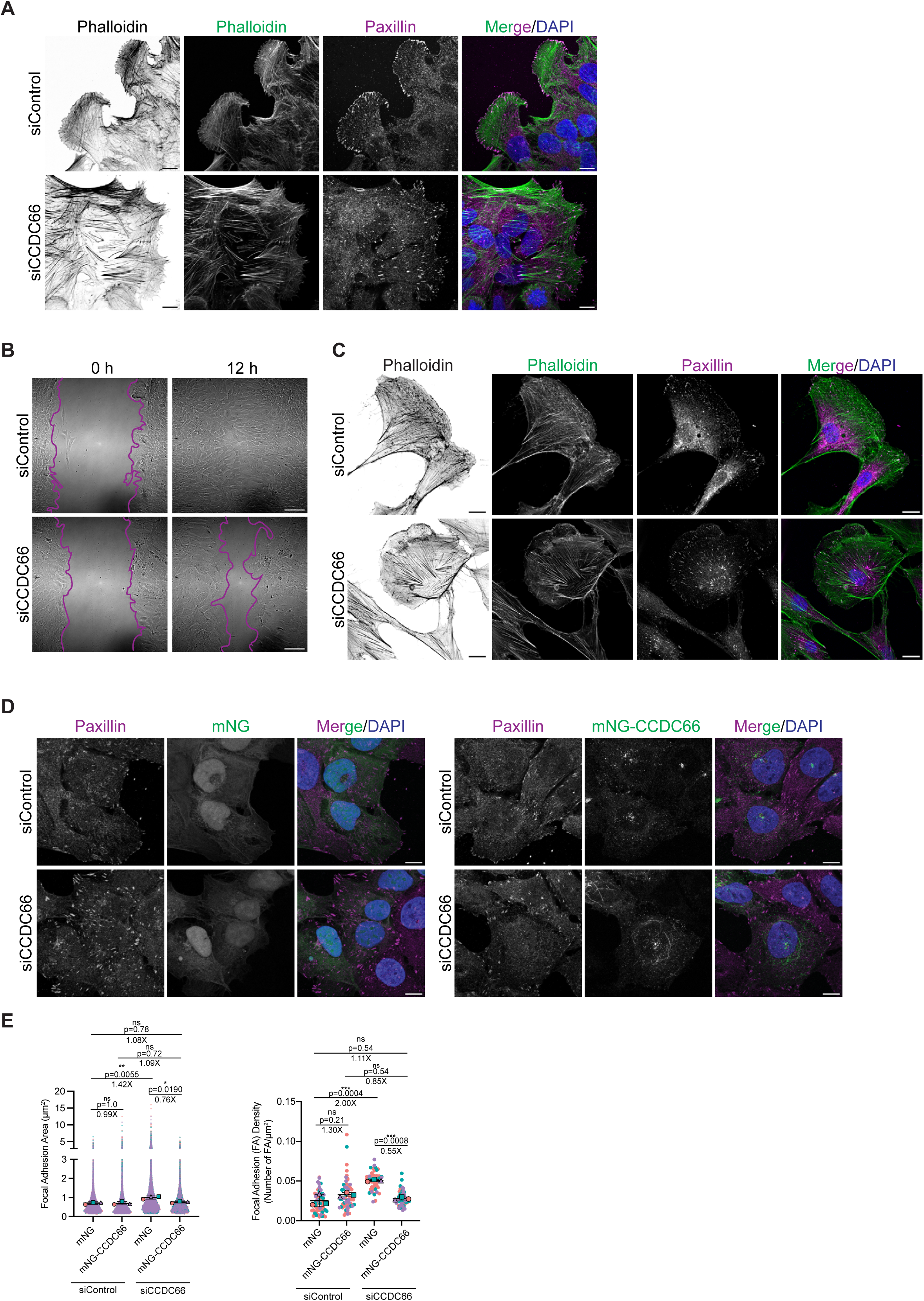
Cell-line validation of CCDC66 depletion phenotypes and rescue of focal adhesion defects. **(A)** Representative immunofluorescence images of U2OS cells at wound edges transfected with control or CCDC66 siRNAs and stained for F-actin with phalloidin, paxillin, and DAPI. Scale bars, 10 µm. **(B)** Representative phase-contrast images of wound healing assays RPE1 cells. Cells were transfected with control or CCDC66 siRNAs. At 72 hours post-transfection, cell monolayers were scratched to create a wound, and images were acquired at 0 h and 12 h post-scratch. Magenta lines outline the migrating wound edges. Scale bars, 100 µm. **(C)** Representative immunofluorescence images of subconfluent RPE1 cells transfected with control or CCDC66 siRNAs and stained for F-actin with phalloidin, paxillin, and DAPI. Scale bars, 10 µm. **(D)** Representative immunofluorescence images of U2OS cells expressing mNG or siRNA-resistant mNG-CCDC66 and transfected with control or CCDC66 siRNAs. Cells were fixed 48 h post-transfection and stained for paxillin to visualize focal adhesions. Nuclei were counterstained with DAPI. mNG fluorescence shows expression of the indicated constructs. Scale bars, 10 µm. **(E)** Quantification of focal adhesion area and focal adhesion density from the experiment shown in (D). Focal adhesion area was calculated from paxillin-positive adhesions, and focal adhesion density was calculated as the number of focal adhesions normalized to cell area. Data are presented as mean ± S.E.M. from *n = 3* independent experiments. Colored dots represent individual measurements, with each color corresponding to one independent experiment. Fold changes and exact p-values are indicated above the graph. Asterisks denote statistical significance (*p < 0.05, **p < 0.01, ***p < 0.001, ns: not significant; two-way ANOVA with Tukey’s post hoc test).

**Figure S3.**
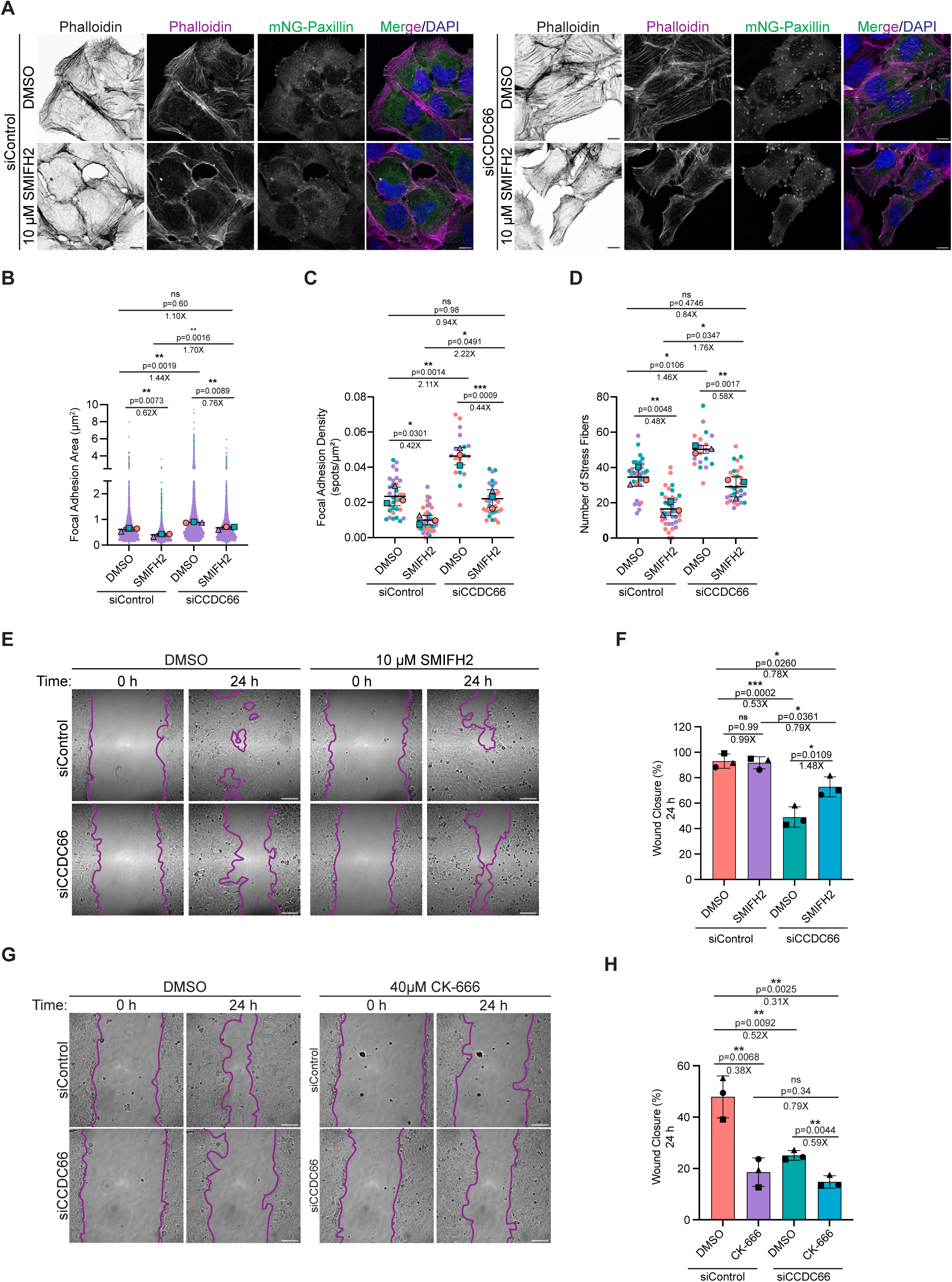
Formin inhibition suppresses stress fiber and focal adhesion accumulation and restores migration in CCDC66-depleted cells. **(A)** Representative immunofluorescence images of U2OS cells stably expressing mNG-Paxillin (green), transfected with control or CCDC66 siRNAs. At 48 h post-transfection, cells were treated with DMSO or SMIFH2 (10 µM) for 1 h, then fixed and stained with phalloidin to visualize F-actin (magenta). Nuclei were counterstained with DAPI (blue). Scale bars, 10 µm. **(B, C)** Quantification of focal adhesion morphology from the images shown in (A), including (B) focal adhesion area (µm^2^), and (C) focal adhesion density (number of FAs per µm^2^ of cell area). Data are mean ± SEM from *n = 3* independent experiments. Colored dots represent individual measurements, with each color corresponding to one independent experiment. Fold changes and exact p-values are indicated above the graph. Asterisks denote statistical significance (*p < 0.05, **p < 0.01, ***p < 0.001, ns: not significant; two-way ANOVA with Tukey’s post hoc test). **(D)** Quantification of the number of stress fibers per cell based on the F-actin staining shown in (A). Data represent mean ± S.E.M. from *n = 3* independent experiments. Colored dots represent individual measurements, with each color corresponding to one independent experiment. Fold changes and exact p-values are indicated above the graph. Asterisks denote statistical significance (*p < 0.05, **p < 0.01; two-way ANOVA with Tukey’s post hoc test). **(E)** Representative phase-contrast images of a wound healing assay in U2OS cells transfected with control or CCDC66 siRNAs at 0 and 24 hours. At 48 h post-transfection, confluent cell monolayers were scratched and treated with DMSO or the formin inhibitor SMIFH2 (10 µM), and imaged every 2 h over a 24-h period. Purple lines outline the migrating wound edges. Scale bars, 100 µm. **(F)** Quantification of wound closure from the assay shown in (E). Wound closure was calculated as the percentage reduction in wound area relative to the initial wound area at 0 h. Data are presented as mean ± SEM from *n = 3* independent experiments. Dots represent independent experiments. Fold changes and exact p-values are indicated above the graph. Asterisks denote statistical significance (*p < 0.05, ***p < 0.001, ns: not significant; two-way ANOVA with Tukey’s post hoc test). **(G)** Representative phase-contrast images of wound healing assays in U2OS cells transfected with control or CCDC66 siRNAs at 0 and 24 hours. At 48 h post-transfection, confluent monolayers were scratched and treated with DMSO or the Arp2/3 inhibitor CK-666 (40 µM), and imaged every 2 h over a 24-h period. Purple lines outline the migrating wound edges. Scale bars, 100 µm. **(H)** Quantification of wound closure from the assay shown in (G). Wound closure was calculated as the percentage reduction in wound area relative to the initial wound area at 0 h. Data are presented as mean ± SEM from *n = 3* independent experiments. Dots represent independent experiments. Fold changes and exact p-values are indicated above the graph. Asterisks denote statistical significance (**p < 0.01, ns: not significant; two-way ANOVA with Tukey’s post hoc test).

**Figure S4.**
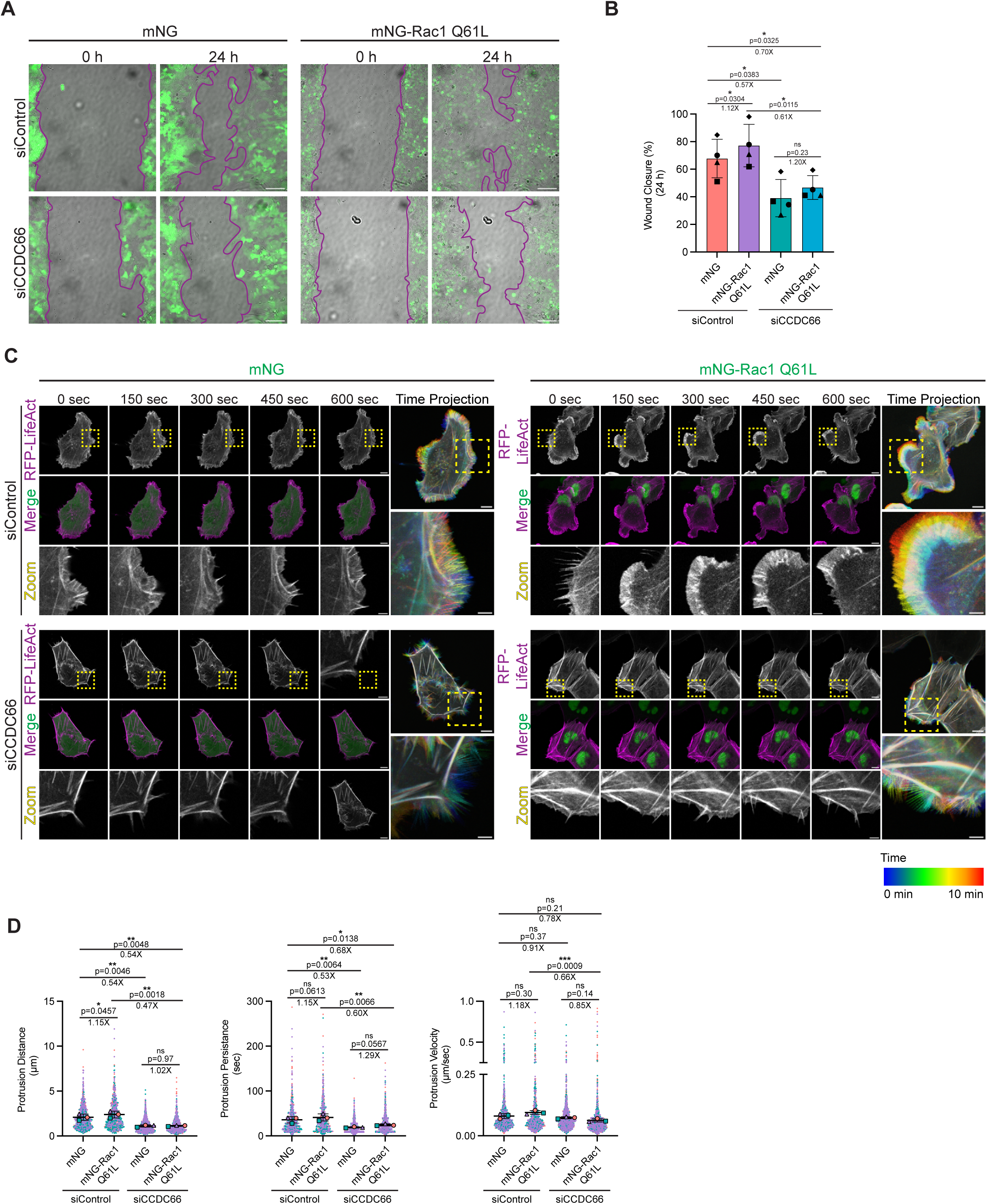
Constitutively active Rac1 does not restore migration or lamellipodial dynamics in CCDC66-depleted cells. **(A)** Representative phase-contrast and fluorescence overlay images from wound-healing assays in U2OS cells stably expressing mNG or constitutively active mNG-Rac1 Q61L. Cells were transfected with control or CCDC66 siRNAs, and confluent monolayers were scratched and imaged at 0 and 24 h post-wounding. Purple lines outline the migrating wound edges; green fluorescence indicates expression of mNG or mNG-Rac1 Q61L. Scale bars, 100 µm. **(B)** Quantification of wound closure from the assay shown in (A). Wound closure was calculated as the percentage reduction in wound area relative to the initial wound area at 0 h. Data are presented as mean ± S.E.M. from *n = 3* independent experiments. Dots represent independent experiments. Fold changes and exact p values are indicated above the graph. Asterisks denote statistical significance (*p < 0.05, ns = not significant; two-way ANOVA with Tukey’s post hoc test). **(C)** Live imaging of U2OS cells expressing RFP-LifeAct and either mNG or mNG-Rac1 Q61L, transfected with control or CCDC66 siRNAs. Cells were imaged every 10 s for 10 min, and representative frames are shown at 0, 150, 300, 450, and 600 s. RFP-LifeAct is shown in grayscale or magenta, and mNG or mNG-Rac1 Q61L is shown in green. Yellow dashed boxes indicate the regions shown at higher magnification in the zoom panels. Temporal color-coded projections are shown to visualize edge dynamics over the imaging period. Scale bars, 10 µm for full-cell images and 3 µm for zoomed regions. **(D)** Quantification of protrusion dynamics from the time-lapse sequences shown in (C), including protrusion distance (µm), protrusion persistence (s), and protrusion velocity (µm/s). Data represent mean ± S.E.M. from *n = 3* independent experiments. Colored dots represent individual measurements, with each color corresponding to one independent experiment. Fold changes and exact p values are indicated above the graphs. Asterisks denote statistical significance (*p < 0.05, **p < 0.01, ***p < 0.001, ns = not significant; two-way ANOVA with Tukey’s post hoc test).

**Figure S5.**
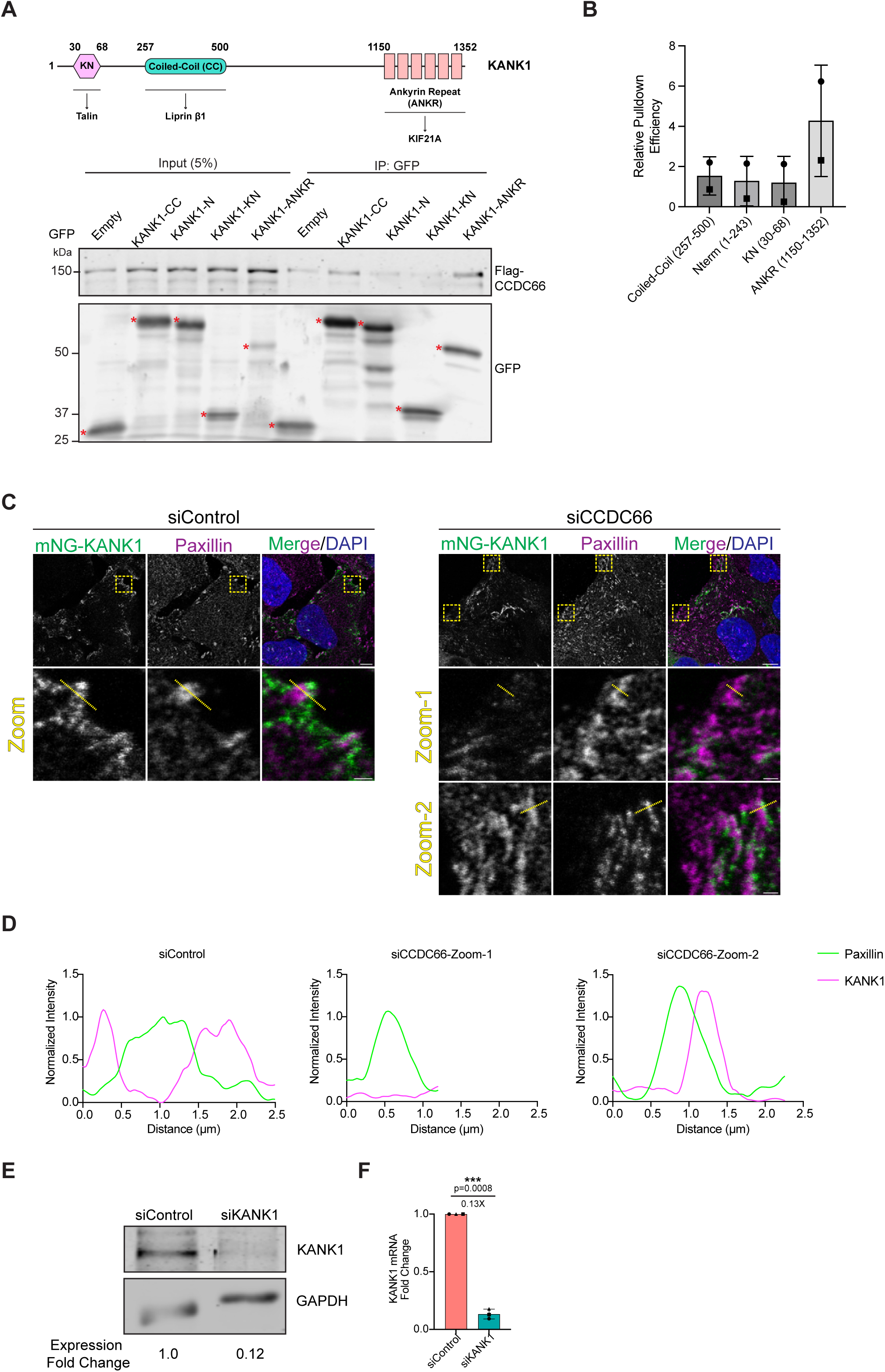
Complementary analyses of the CCDC66-KANK1 interaction, KANK1 redistribution, and validation of KANK1 depletion. **(A)** Domain mapping of the CCDC66-KANK1 interaction. Top, schematic of KANK1 domain organization and GFP-tagged truncation constructs used for pulldown analysis, including the coiled-coil region (CC, aa 257-500), N-terminus (N, aa 1-243), KN motif (KN, aa 30-68), and ankyrin-repeat region (ANKR, aa 1150-1352). Known interaction regions for talin, liprin-β1, and KIF21A are indicated. Bottom, HEK293T cells were co-transfected with FLAG-CCDC66 and either GFP alone or the indicated GFP-tagged KANK1 fragments. Cell lysates were subjected to GFP-trap pulldown and analyzed by immunoblotting with anti-FLAG and anti-GFP antibodies. Input represents 5% of total cell lysate. Red asterisks indicate the expected GFP-tagged bait proteins, and the relative pulldown efficiency calculated from the representative blot is shown on the right (B). **(B)** Quantification of relative FLAG-CCDC66 pulldown efficiency for the KANK1 fragments shown in (A). Relative pulldown efficiency was calculated as the ratio of co-precipitated FLAG-CCDC66 in the GFP IP, normalized to the corresponding FLAG-CCDC66 input and recovered GFP bait, and then normalized to the GFP control. Data are presented as mean ± S.E.M. from *n = 2* independent experiments. Symbols represent independent experiments. **(C)** Representative confocal images of U2OS cells expressing mNG-KANK1 and transfected with control or CCDC66 siRNAs. Cells were stained for paxillin to mark focal adhesions, and nuclei were counterstained with DAPI. Yellow dashed boxes indicate peripheral regions shown at higher magnification below, and yellow dashed lines indicate the paths used for line-scan analysis in (D). Two siCCDC66 zoom regions are shown to illustrate disrupted KANK1 organization and redistribution into broader or fiber-like peripheral structures. Scale bars, 10 μm for full-cell images and 1 μm for magnified regions. **(D)** Representative line-scan intensity profiles of mNG-KANK1 and paxillin across the regions marked in (C). Fluorescence intensities across all channels and experimental conditions were normalized to the maximum Paxillin signal in the siControl group. Control cells show mNG-KANK1 enrichment adjacent to paxillin-positive adhesions. The two representative siCCDC66 profiles illustrate either reduced KANK1 enrichment at the adhesion edge or displacement/redistribution of KANK1 relative to the paxillin-positive adhesion core. **(E)** Immunoblot validation of KANK1 depletion at the protein level. U2OS cells were transfected with control or KANK1 siRNAs, and whole-cell lysates were collected 48 h post-transfection. KANK1 protein levels were analyzed by immunoblotting, with GAPDH used as a loading control. **(F)** Validation of KANK1 knockdown at the mRNA level. U2OS cells were transfected with control or KANK1 siRNAs, and total RNA was collected 48 h post-transfection for qRT-PCR analysis. KANK1 transcript levels were normalized to GAPDH. Data represent mean ± S.E.M. from *n = 3* independent experiments. Dots represent independent experiments. Fold change relative to siControl is indicated above the graph. Asterisks denote statistical significance (***p < 0.001; paired Student’s t-test).

**Figure S6.**
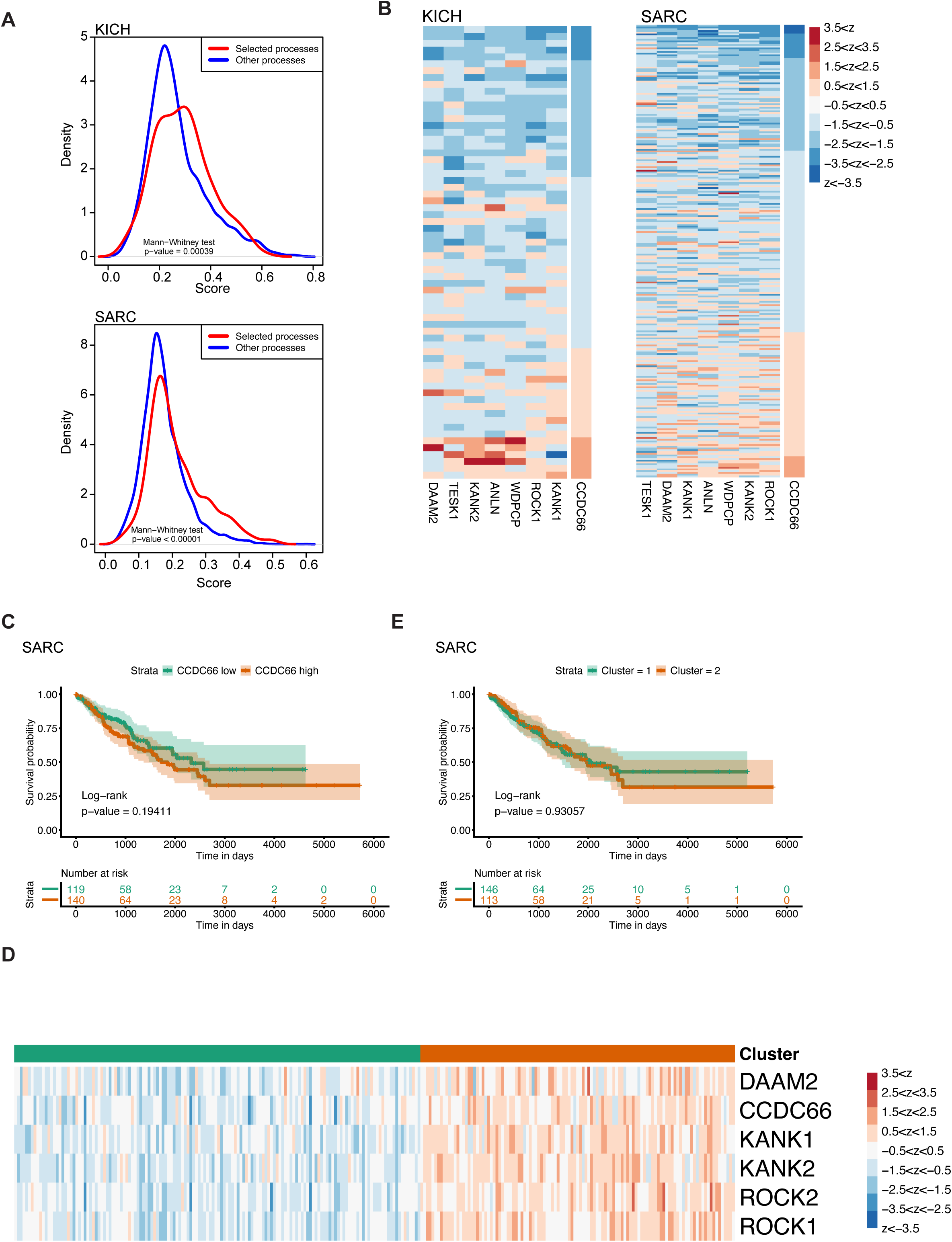
CCDC66-associated migration and cytoskeletal programs are recurrent across tumor cohorts and show context-dependent survival associations. **(A)** Distribution of GO Biological Process enrichment scores for predefined migration-, adhesion-, cytoskeleton-, and microtubule-associated GO categories compared with all other GO terms in KICH and SARC. GO enrichment analysis was performed using genes correlated with CCDC66 expression. Red curves indicate the selected functional categories, and blue curves indicate all other GO terms. Statistical significance was determined by Mann-Whitney test, with exact p values indicated in the plots. **(B)** Heatmaps of z-scored expression values for genes within the GO term podocyte cell migration in SARC (left) and KICH (right) tumors. These enriched gene sets include CCDC66 together with multiple genes linked to the cytoskeletal program defined in this study, including KANK1, KANK2, ROCK1, and DAAM2. **(C)** Kaplan-Meier analysis of overall survival in SARC patients stratified into CCDC66-low and CCDC66-high groups. CCDC66 expression alone did not significantly stratify overall survival. The number of patients in each group is indicated in the plot. P values were calculated by log-rank test. **(D)** Heatmap of z-scored expression values for the CCDC66/KANK/ROCK-associated module (DAAM2, CCDC66, ROCK2, KANK1, KANK2, and ROCK1) in SARC tumors grouped by unsupervised clustering. The top annotation bar indicates the two clusters. **(E)** Kaplan-Meier analysis of overall survival for the two SARC clusters shown in (D). Grouping tumors by the broader CCDC66/KANK/ROCK-associated module did not significantly stratify overall survival, indicating that the clinical association of this cytoskeletal program is context dependent. The number of patients in each group is indicated in the plot. P values were calculated by log-rank test.

